# Defining Dementia Subtypes Through Neuropsychiatric Symptom-Linked Brain Connectivity Patterns

**DOI:** 10.1101/2023.07.02.547427

**Authors:** Kanhao Zhao, Hua Xie, Gregory A. Fonzo, Nancy Carlisle, Ricardo S. Osorio, Yu Zhang

**Author notes:** Corresponding author: Yu Zhang, Assistant Professor of Bioengineering, Assistant Professor of Electrical and Computer Engineering, Lehigh University, Bethlehem, PA 18015, USA.

## Abstract

**BACKGROUND:** Dementia is highly heterogeneous, with pronounced individual differences in neuropsychiatric symptoms (NPS) and neuroimaging findings. Understanding the heterogeneity of NPS and associated brain abnormalities is essential for effective management and treatment of dementia.

**METHODS:** Using large-scale neuroimaging data from the Open Access Series of Imaging Studies (OASIS-3), we conducted a multivariate sparse canonical correlation analysis to identify functional connectivity-informed symptom dimensions. Subsequently, we performed a clustering analysis on the obtained latent connectivity profiles to reveal neurophysiological subtypes and examined differences in abnormal connectivity and phenotypic profiles between subtypes.

**RESULTS:** We identified two reliable neuropsychiatric subsyndromes – behavioral and anxiety in the connectivity-NPS linked latent space. The behavioral subsyndrome was characterized by the connections predominantly involving the default mode and somatomotor networks and neuropsychiatric symptoms involving nighttime behavior disturbance, agitation, and apathy. The anxiety subsyndrome was mainly contributed by connections involving the visual network and the anxiety neuropsychiatric symptom. By clustering individuals along these two subsyndromes-linked connectivity latent features, we uncovered three subtypes encompassing both dementia patients and healthy controls. Dementia in one subtype exhibited similar brain connectivity and cognitive-behavior patterns to healthy individuals. However, dementia in the other two subtypes showed different dysfunctional connectivity profiles involving the default mode, frontoparietal control, somatomotor, and ventral attention networks, compared to healthy individuals. These dysfunctional connectivity patterns were associated with differences in baseline dementia severity and longitudinal progression of cognitive impairment and behavioral dysfunction.

**CONCLUSIONS:** Our findings shed valuable insights into disentangling the neuropsychiatric and brain functional heterogeneity of dementia, offering a promising avenue to improve clinical management and facilitate the development of timely and targeted interventions for dementia patients.

## Introduction

Dementia is highly heterogenous, as reflected by variability in genetic risk factors, neuropsychiatric symptoms (NPS), neuroimaging, comorbidities and copathology (1, 2). Conventional analyses of dementia using neuroimaging and psychological behavior have focused on characterizing case-control group differences, assuming homogeneity among dementia patients (3–5). However, this approach neglects the inherent heterogeneity of dementia NPS and brain abnormalities, limiting our understanding of the underlying mechanisms of dementia.

To untangle the heterogeneity of NPS of dementia, some studies focus on analyzing NPS subsyndromes, as it has been proposed that some NPS can be clustered into several neuropsychiatric subsyndromes (6). These subsyndromes often co-occur and show a similar clustering pattern of NPS, and provide insights into the pathophysiology of various cognitive impairments seen in different types of dementia (7) and the effectiveness of some dementia drug treatments (8). For instance, neuroleptic interventions have been shown to alleviate symptoms in some neuropsychiatric subsyndromes rather than specific NPS, e.g. delusions from other behavioral manifestations (9, 10). However, the traditional subsyndrome analytic framework, which used matrix decomposition of NPS score to obtain the subsyndromes (7, 11, 12) and then compared their differences across various dementia categories (4, 13), has two major limitations. Firstly, the purely unsupervised matrix decomposition of NPS fails to provide a clear neurobiological basis for the subsyndromes and shows a clear link to corresponding heterogeneous brain abnormalities. Secondly, typical dementia diagnostic categories derived from molecular phenotypes (14) may not adequately reflect the heterogeneity of NPS and alight well with underlying neurobiology observed in dementia patients (4).

In recent years, researchers have employed data-driven multivariate methods to establish connections between neuroimaging features and psychological assessments for disentangling the shared heterogeneity across these two modalities (15–18). Two studies have utilized sparse canonical correlation analysis (sCCA) or partial least square analysis to uncover latent subsyndromes by linking phenotypes with functional connectivity (FC), which quantifies synchronization in resting-state functional resonance magnetic imaging (fMRI) signals among different brain regions (15, 16). Using a similar method to derive subsyndromes for depression or autism, two additional studies further defined subtypes to parse the neurobiological heterogeneity present within these conditions (17, 18). However, the lack of cross-validation limits the robustness and generalizability of the identified subsyndromes. More critically, no study has successfully derived FC-informed neuropsychiatric subsyndromes or identified potential subtypes by utilizing latent connectivity-NPS linked features, thus limiting our understanding of the heterogeneity within dementia.

In this study, we sought to employ sCCA (19) to identify cross-validated FC-informed symptom dimensions (subsyndromes) across dementia and healthy individuals exhibiting at least one neuropsychiatric symptom. Furthermore, we aimed to reveal stable subtypes for those subjects based on the identified subsyndromes. By accomplishing these goals, we expect to enhance our comprehension of the relationship between functional connectivity and NPS, as well as to elucidate the heterogeneity observed in dementia. Specifically, utilizing the Open Access Series of Imaging Studies (OASIS-3) dataset, we linked FC and NPS in the latent space using sCCA. This analysis resulted in the identification of two distinct neuropsychiatric subsyndromes: the behavioral subsyndrome and the anxiety subsyndrome. The behavioral subsyndrome was characterized by prominent connections involving the default mode network (DMN) and somatomotor network (SMN), and the neuropsychiatric symptoms in nighttime behavior disturbance, agitation, and apathy. On the other hand, the anxiety subsyndrome was characterized by connections involving the visual network (VIS) and the presence of anxiety symptom. Statistical analyses further confirmed associations between these two subsyndromes and other phenotypic characteristics, including functional disability. Furthermore, by employing K-means clustering, we identified three distinct and stable subtypes based on FC features linked to the subsyndromes. To gain deeper insight into the psychological underpinnings of dementias of these subtypes, we compared NPS, FCs, and the severity of cognitive-behavioral dysfunction at baseline across dementias of the three subtypes. Additionally, we explored the long-term progression of cognitive-behavioral dysfunction within these subtypes. The proposed analytical framework for this study is illustrated in Figure 1.

**Figure 1.**
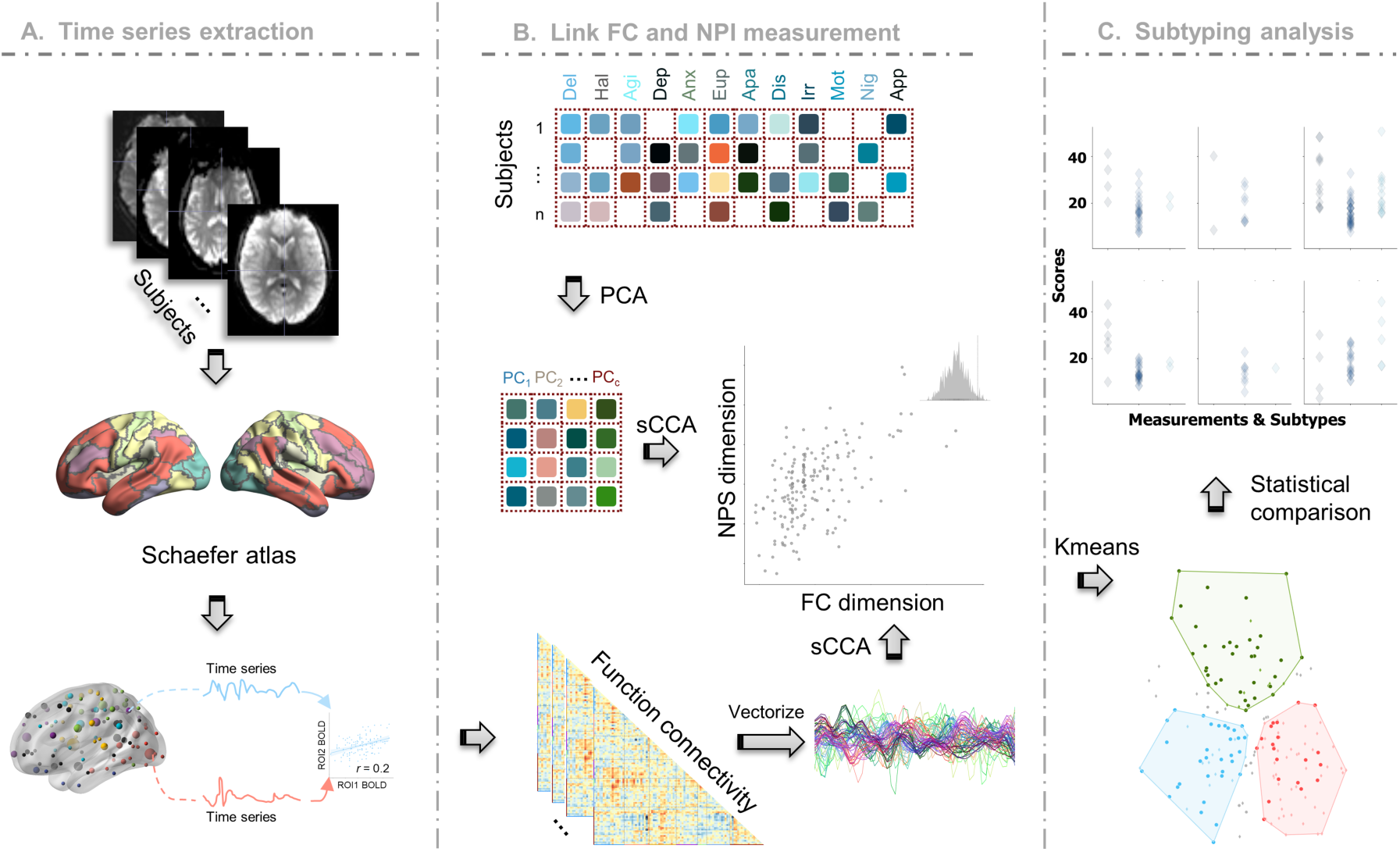
Framework of our study. **A** The BOLD signals extracted from the preprocessed fMRI, were grouped into 100 regions of interests (ROIs) defined by the Schaefer parcellation. FCs were obtained by calculating the Pearson correlation between pairwise ROI time signals. **B** PCA was applied to decompose the NPS into low-dimensional orthogonal latent space. sCCA was then employed to maximize the correlation between FCs and principal components of NPS profile. **C** Subtype identification was achieved through Kmeans clustering along the NPS subsyndrome-linked FC latent features.

## METHODS

### Participants

In this study, we utilized the longitudinal data collected from the Open Access Series of Imaging Studies (OASIS-3) database, which enrolled participants at Washington University in St. Louis over a period of 15 years (20). The dataset comprised resting-state fMRI scans and clinical assessments as well as neuropsychological measures from 1098 participants aged between 42 to 95. Exclusion criteria included medical conditions that precluded longitudinal participation (e.g. end-stage renal disease requiring dialysis) or medical contraindications for the study arms (e.g. pacemaker for MRI, anticoagulant use for lumbar puncture).

### Neuropsychiatric Inventory

The *Neuropsychiatric Inventory* (21) is a commonly used measure for evaluating neuropsychiatric related behavioral and psychological symptoms. It consists of twelve domains: delusions, hallucinations, agitation, depression, anxiety, euphoria, apathy, disinhibition, irritability, aberrant motor behavior, nighttime behavior disturbance, and appetite abnormality. The neuropsychiatric inventory assesses symptoms in four levels: 0 (no symptom), 1 (mild but not significant change), 2 (significant but not dramatic change), 3 (dramatic change).

### Other clinical and psychological assessments

To assess functional impairment, we utilized multiple measures *Clinical Dementia Rating Scale (CDR), Mini-Mental State Examination (MMSE), and Functional Activities Questionnaire (FAQ).* The CDR evaluates cognitive impairment in six domains including memory, orientation, judgment and problem solving, community affair, home and hobbies and personal care, and provides a global CDR score and the sum of boxes (SOB) (22). The CDR range from 0–3: no dementia (CDR = 0), questionable dementia (CDR = 0.5), MCI (CDR = 1), moderate cognitive impairment (CDR = 2), and severe cognitive impairment (CDR = 3). The MMSE is also designed for the evaluation of cognitive impairment (23), but only the total score was accessible. Smaller MMSE denotes more severe cognitive dysfunction. The functional ability is evaluated by FAQ (24) in 10 daily activities: 1) writing checks, paying bills, and keeping financial records; 2) assembling tax records and making out business and insurance papers; 3) shopping alone for clothes, household necessities, and groceries; 4) playing a game of skill such as bridge, other card game, or chess; 5) heating water for coffee or tea and turning off the stove; 6) preparing a balanced meal; 7) keeping track of current events; 8) paying attention to and understanding a TV program, book, or magazine; 9) remembering appointments, family occasions, and medications; and 10) travel out of the neighborhood. Moreover, the *neuropsychological assessment battery (NAB)* (25) was applied in assessing neuropsychiatric disturbances and cognitive and behavioral dysfunction. For the NAB, ten neuropsychological tests measure attention/working and episodic memory, executive function, and language, including the Digit Span Forward and Backward test, Logical Memory, and Digit Symbol Coding in Wechsler Memory Scale (WAIS-R) (26), Category fluency of animal and vegetable (FLU-ANI/ FLU-VEG) (27), Trail Making Test Part A and B (28), and Boston Naming Test (29).

### Image acquisition and preprocessing

Neuroimaging data in OASIS-3 was scanned in 3 different Siemens scanners (Siemens Vision 1.5 T, 2 scanners of TIM Trio 3T) and Siemens BioGraph mMR PET-MR 3T. High resolution T1-weighted structural image (TR = 2.4 s, TE = 3.08 ms, FOV = 256 × 256 mm, FA = 8°, voxel size 1 × 1 × 1 mm^3^) and resting-state functional image (EPI; TR = 2.2 s, TE = 27 ms, FOV = 240 × 240 mm, FA = 90°, duration = 6 min, voxel size 4 × 4 × 4 mm, 36 slices) were used. This study focused on the analysis of participants who had both fMRI and at least one neuropsychiatric symptom at baseline. As a result, a total of 177 subjects were utilized.

The acquired rs-fMRI data were preprocessed using the reproducible fMRIPrep pipeline (30). The T1 weighted image was corrected for intensity nonuniformity and then skull stripped. Spatial normalization was done through nonlinear registration, with the T1w reference (31). Using FSL, brain tissue such as cerebrospinal fluid, white matter, and grey matter was segmented from the reference, brain-extracted T1 weighted image (32). The fieldmap information was used to correct distortion in low-frequency and high-frequency components of fieldmap caused by field inhomogeneity. With less fieldmap distortion, a corrected echo-planar imaging reference was obtained from a more accurate co-registration with the anatomical reference. The blood oxygenation level dependent (BOLD) reference was then transformed to the T1-weighted image with a boundary-based registration method, configured with nine degrees of freedom to account for distortion remaining in the BOLD reference (33). Head-motion parameters (rotation and translation parameters of volume-to-reference transform matrices) were estimated with MCFLIRT (FSL). BOLD signals were slice-time corrected and resampled onto the participant’s original space with head-motion parameters, susceptibility distortion correction, and then resampled into standard space (MNI152NLin2009cAsym space), generating a preprocessed BOLD signal. Automatic removal of motion artifacts using independent component analysis (ICA-AROMA) (34) was performed on the preprocessed BOLD time-series in MNI space after removal of non-steady-state volumes and spatial smoothing with an isotropic Gaussian kernel of 6 mm FWHM (full-width half-maximum).

### Calculation of resting-state functional connectivity

The voxel time series were averaged into time series of 100 regions of interest (ROIs) defined by the Schaefer parcellation (35). Pearson correlation was then computed between time series of each pair of ROIs, resulting in 4950 FCs for each participant. Fisher’s r-to-z transformation was applied to enhance normality of connectivity, followed by z-score normalization.

### Connectivity-NPS linked dimension analysis

We sought to identify latent dimensions linking functional connectivity and NPS in a data-driven manner. The analysis details are described below.

### Dimensionality reduction

Significant correlations among NPS symptoms were observed in previous study (1) as well as in our study (Figure S1). The presence of collinearity among NPS symptoms may result in ill-conditioning for the subsequent canonical correlation analysis step (36). To address this issue, we employed principal component analysis (PCA) to decompose high-dimensional collinear features into orthonormal bases. Seven orthogonal principal components extracted from twelve NPS domains (Figure S2A, >80% variance explained) were used for further sparse canonical correlation analysis.

### Sparse canonical correlation analysis

We employed sparse canonical correlation analysis (sCCA) to identify canonical variates that exhibited maximum correlation between the projected principal components of NPS and FC across both healthy controls and patients. As a multivariate statistical approach, sCCA identifies linear transformations that result in maximal correlation between matched canonical variates while alleviating the overfitting issues through sparse constraints, achieved by L1 regularization (37). Model performance was assessed by calculating the average correlation of pairwise canonical variates in ten-fold cross validation. To determine the optimal hyperparameters for L1 regularization and the number of principal components, we conducted a grid search within the inner loop of ten-fold cross-validation, aiming to maximize the average correlation of available canonical variates. As indicated in Figure S2B, the highest average correlation was obtained when using seven principal components and an L1 regularization penalty of 0.6. For subsequent analyses, we focused on the first and second canonical variates since they accounted for nearly 80% covariance between FC and NPS (Figure S2C). Importantly, our cross-validation showed that only these two canonical FC variates exhibited significant correlation with their corresponding canonical NPS variates and survived the permutation test with the false discovery rate (FDR) correction of cross-validated correlation coefficients of all decomposed canonical variates (Figures 2A, 3A, Table S1).

**Figure 2.**
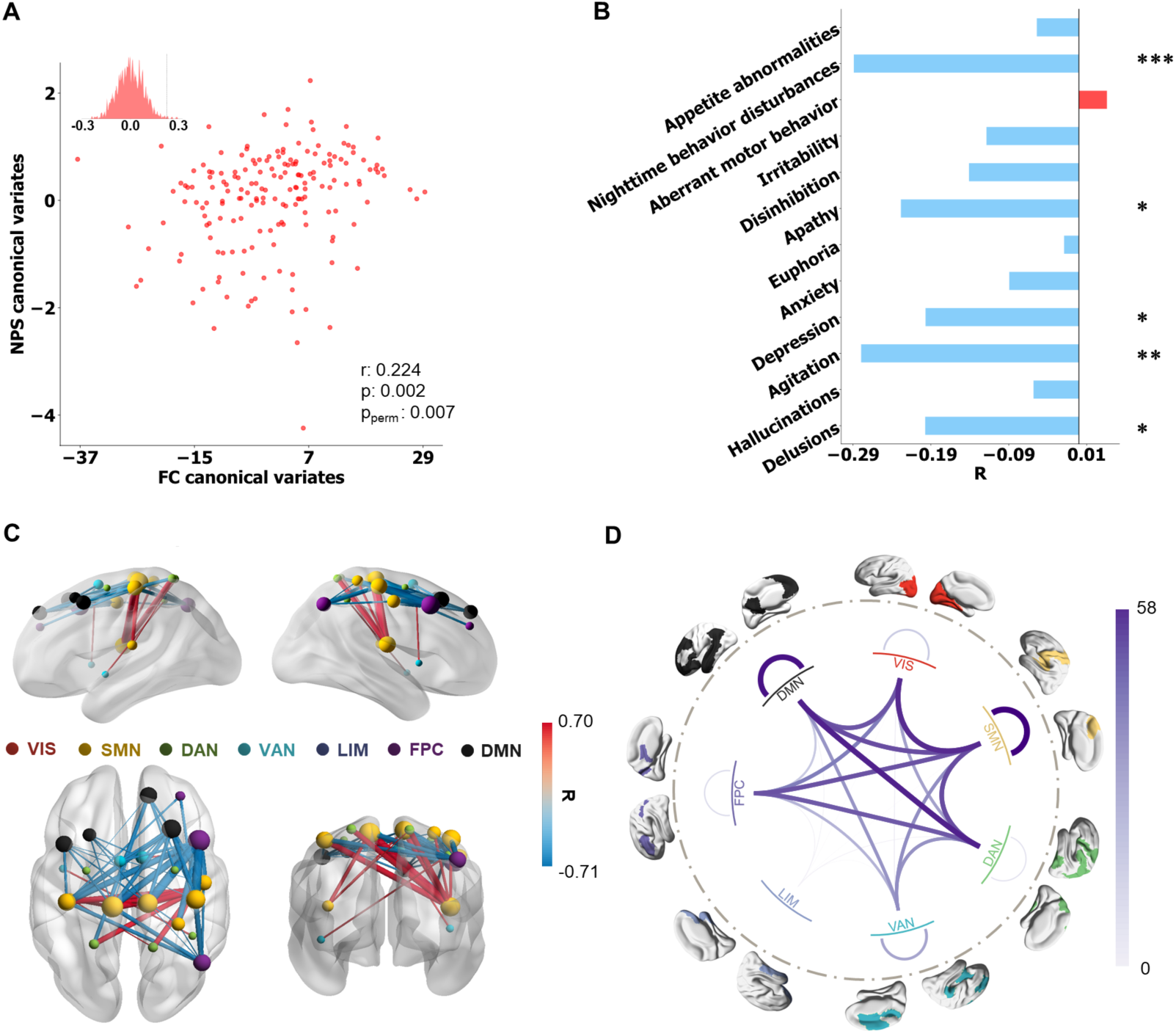
Behavioral subsyndrome profile and linked FC pattern. **A** Correlation between FC and NPS latent features, evaluated by 10-fold cross-validation (r = 0.22, p = 0.002 and p_permute_ = 0.007). **B** Correlation coefficients between the behavioral subsyndrome-linked FC latent features and NPS. The statistical significance of correlation is denoted by *(p ≤ 0.05), **(p ≤ 0.01), ***(p ≤ 0.001), and ****(p ≤ 0.0001). **C** Top 50 (absolute) strongest correlations (FDR correction) between the behavioral subsyndrome-linked FC latent features and FC features. The FCs are depicted in red or blue based on their positive or negative correlation, respectively. Node size indicated the node strength calculated from the summed FC strength of each ROI. **D** All significant (p_fdr_ < 0.05) correlation coefficients between the behavioral subsyndrome-linked FC latent features and FC features were grouped into Yeo’s 7 networks, including visual network (VIS), somatomotor network (SMN), dorsal attention network (DAN), ventral attention network (VAN), limbic network (LIM), frontoparietal control network (FPC), default mode network (DMN).

### Clustering-based identification of neurophysiological subtypes

Using the neuropsychiatric subsyndrome-linked latent FC features, we employed the K-means clustering approach to identifying distinct neurophysiological subtypes in dementia. Considering the importance of incorporating healthy controls alongside dementia patients to capture the heterogeneity of dementias, we included both groups in the clustering analysis, drawing inspiration from the concept of normative modeling, which utilizes the healthy group as a reference (38). To evaluate the clustering performance, we calculated Calinski Harabasz and silhouette metrics by conducting 1000 trials of randomly subsampling 90% of the subjects (39, 40). Both metrics serve as internal cluster validation indices, with higher values indicating better clustering. The Calinski Harabasz metric quantifies the ratio of between-cluster variance to within-cluster variance, while the silhouette metric calculates the ratio of difference between the intra-cluster distance and nearest-cluster distance to difference between the maximum of the intra-cluster distance and nearest-cluster distance. Additionally, we assessed the stability of cluster assignments by testing whether pairs of subjects were consistently assigned to the same cluster across 1000 random subsamplings. To confirm the robustness of our subtype findings in relation to the selection of clustering approach, we performed the same subtyping analysis using hierarchical clustering (Ward’s method) and compared the results with those obtained using K-means clustering.

### Post-hoc analyses

Having identified neuropsychiatric subsyndromes, we further examined their associations with original FCs and NPS, respectively. We calculated the Pearson correlation coefficient to assess the relationship between the FC and subsyndrome-linked FC latent features. Since neuropsychiatric symptoms were ordinal variables, we calculated the Spearman correlation coefficient to evaluate the association between NPS and subsyndrome-linked FC latent features. Additionally, we examined the associations between other continuous phenotypic characteristics (such as age and MMSE) and subsyndromes-linked FC latent features using Pearson correlation. For discrete phenotypic characteristics, such as gender and presence of functional behavioral disability, we used analysis of variance to evaluate the associations with subsyndrome-linked FC latent features.

To investigate neural circuit abnormalities within each subtype, we conducted a comparative analysis of FC between dementia patients in each subtype and healthy controls using the Wilcoxon signed-rank test. To account for multiple comparisons, we applied FDR to correct the p-values of the detected FC differences for each subtype. Additionally, we utilized the Chi-square test to identify associations between categorical phenotypes and dementia patients in each subtype. For ordinal phenotypes, we employed the Kruskal-Wallis analysis of variance to detect differences across dementia patients in each subtype. In cases where significant phenotypic differences were observed across all subtypes, we used Dunn’s test to further examine the pairwise relationships. We corrected the significance of Dunn’s test results for each assessment using FDR. To validate the longitudinal cognitive abnormal progression of dementia within each subtype, we employed linear mixed-effect models. These models included the characteristic assessment score at each study visit as the dependent variable, while dementia subtype label, time state, and the interaction of subtype label and time as independent variables. We applied FDR correction to the p-values of interaction effects in all phenotypic items.

## RESULTS

### Behavioral subsyndrome and linked functional connectivity pattern

The first latent component showed a significant correlation between FC and NPS latent features (r

= 0.22, p = 0.002 and p_permute_ = 0.007) (Figure 2A), as confirmed through 10-fold cross-validation. Following the finding in a previous study (41), we named the first latent component *behavioral subsyndrome* based on its significantly negative correlation with the NPS scores in domains of nighttime behavior disturbance, agitation, and apathy (Figure 2B). To explore its neurobiology, we calculated Pearson correlation between each FC and the behavioral subsyndrome-linked FC latent features. The most significantly correlated FCs were visualized in Figure 2C, while all significant correlations surviving the FDR correction were shown in Figure S3A and grouped according to Yeo’s 7 networks (42) (Figure 2D). The behavioral subsyndrome-linked FCs were characterized by connectivity within SMN and DMN, as well as between-network connectivity involving FPC, SMN, DAN, and DMN. Additionally, we observed that the FC latent features exhibited significantly positive correlations with total scores of MMSE and FLU-VEG while showing negative correlation with CDR-SOB (Table S4). Futhermore, a more severe behavioral subsyndrome was associated with lower scores in CDR domains of memory, home & hobbies, judgment & problem solving, orientation, and community affairs and increased disability in paying attention evaluated by FAQ (Table S3, Figures S4 and S5), indicating that greater behavioral subsyndrome severity was linked to better cognitive performance.

### Anxiety subsyndrome and linked functional connectivity pattern

A significant correlation was also observed for the second latent component between FC and NPS latent features (Figure 3A, r = 0.19, p = 0.01 and p_permute_ = 0.006). This latent component was designated as the anxiety subsyndrome, as it showed the highest positive correlation with the NPS anxiety score (Figure 3B). The FCs that exhibited dominant and significant correlations with the anxiety subsyndrome-linked FC latent features are depicted in Figure 3C (all significantly correlated FCs surviving FDR correction are visualized in Figure S3B). The FC showing the strongest positive correlation with the anxiety subsyndrome was between the right middle frontal cortex (DMN) and supramarginal gyrus (VIS), while the FC between the left superior temporal cortex (SMN) and middle occipital cortex (VIS) exhibited the strongest negative correlation with the anxiety subsyndrome. When grouping the correlations into seven networks,it was observed that the anxiety subsyndrome primarily correlated with connections within the VIS and between-network connections involving VIS, SMN and DMN (Figure 3D). Moreover, the anxiety subsyndrome score exhibited a significant positive correlation with the CDR-SOB score, FAQ paying attention score and negative correlation with the MMSE, FLU-VEG, FLU-ANI, and WAIS-R scores (Tables S3 and S4, Figure S5). A larger anxiety subsyndrome was associated with increased scores across five domains of the CDR, including memory, home & hobbies, judgment & problem solving, orientation, and community affairs (Table S3, Figure S4), indicating that a more pronounced anxiety subsyndrome was associated with a more severe cognitive deficit. Notably, males exhibited significantly larger anxiety subsyndrome scores compared to females (Table S2).

**Figure 3.**
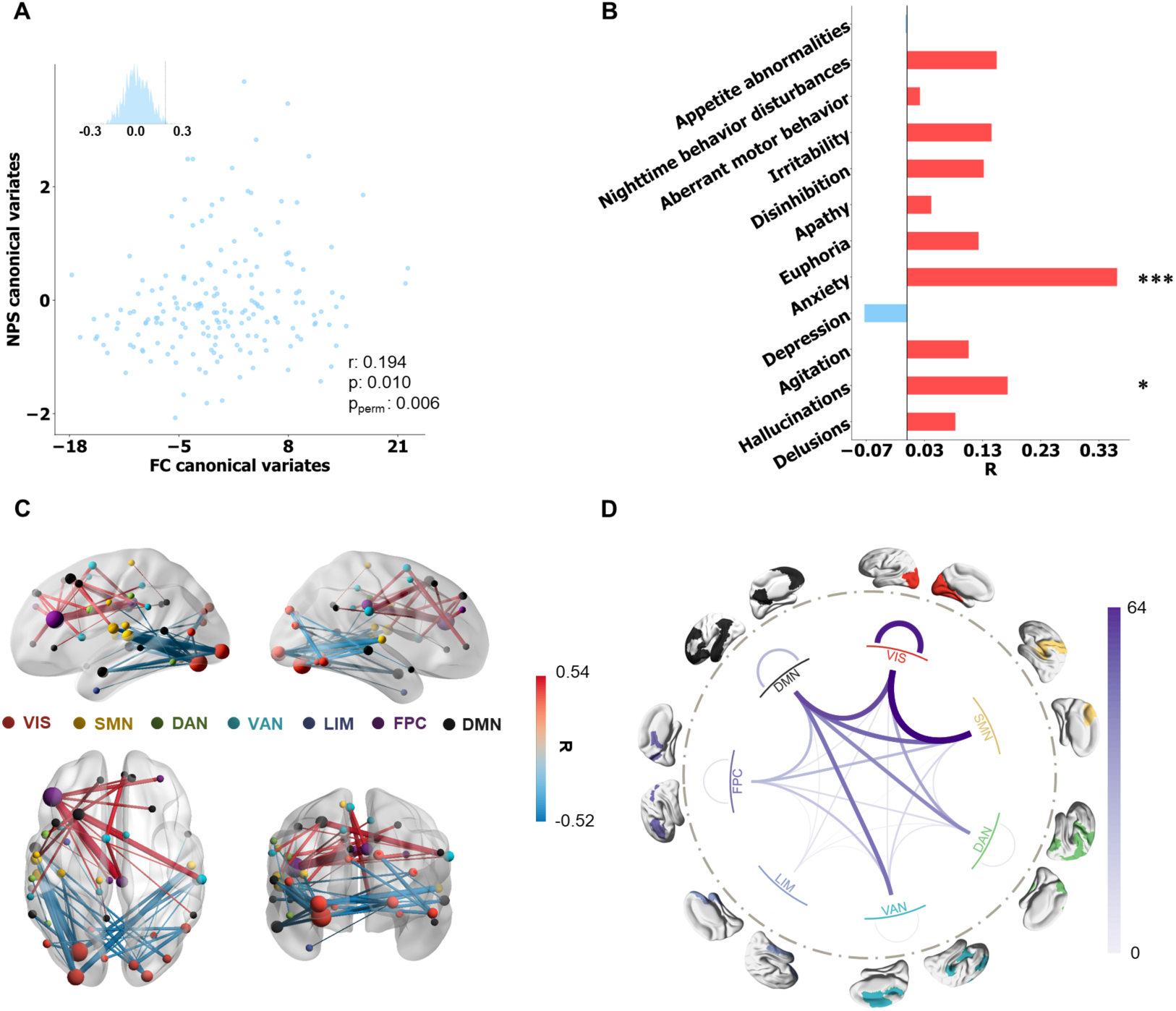
Anxiety subsyndrome profile and linked FC pattern. **A** Correlation between FC and NPS latent features, evaluated by 10-fold cross validation (r = 0.194, p = 0.010 and p_permute_ = 0.006). **B** Correlation coefficients between the anxiety subsyndrome-linked FC latent features and NPS symptoms. The statistical significance of correlation is denoted by *(p ≤ 0.05), **(p ≤ 0.01), ***(p ≤ 0.001), and ****(p ≤ 0.0001). **C** Top 50 (absolute) strongest correlations (FDR correction) between the anxiety subsyndrome-linked FC latent features and FC features. The FCs are depicted in red or blue based on their positive or negative correlation, respectively. Node size indicated the node strength calculated from summed FC strength of each ROI. **D** All significant (p_fdr_ < 0.05) correlation coefficients between the behavioral subsyndrome-linked FC latent features and FC features were grouped into Yeo’s 7 networks.

### Behavioral and anxiety subsyndromes-linked FC latent features define three subtypes

To further disentangle the neurobiological and neuropsychiatric heterogeneity of dementias, we performed K-means clustering analysis on all subjects to reveal subtypes, using anxiety and behavioral subsyndromes-linked FC latent features. We seeked the distinct characteristic profiles of dementia within these subtypes. As a result, we successfully identified three distinct neurophysiological subtypes (Figure 4A). All evaluation metrics of our clustering analysis confirmed the highest level of separability and stability among these three subtypes (Figure S6). Furthremore, the corroborating findings from hierarchical clustering analysis (Figure S7) further demonstrated the robustness of our subtype findings, as verified by a Jaccard similarity of 1 between the derived subtype labels and those from K-means clustering.

**Figure 4.**
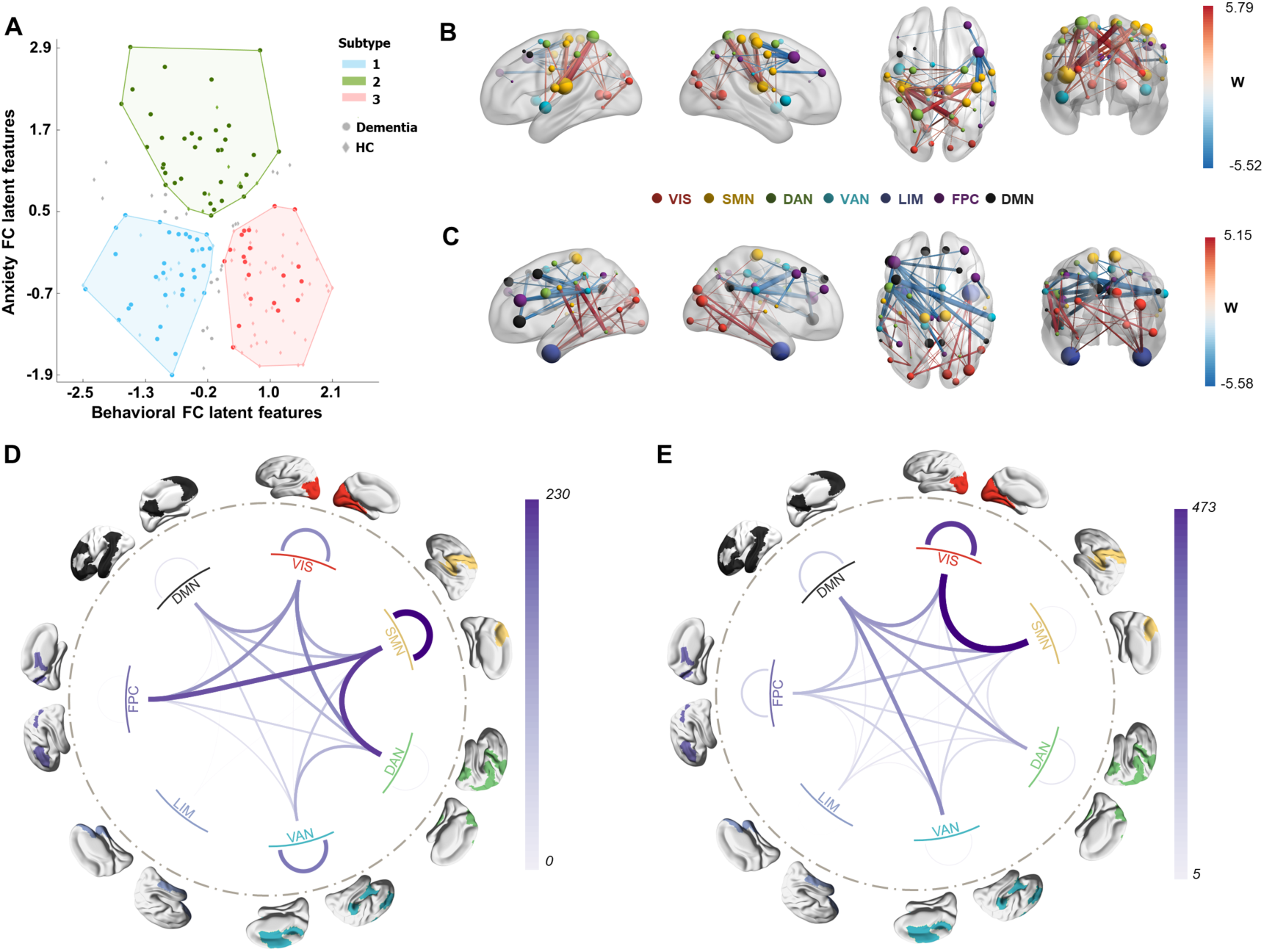
Subtypes defined from NPS subsyndromes-linked FC latent features. **A** Scatter plot of three clusters of subjects along the dimension of anxiety subsyndrome and behavioral subsyndrome. Deeper color indicated dementia patients, while light color indicated healthy controls. Grey dots indicated the subjects who had a greater than 5% chance of being assigned to different subtype labels in 1000 random 90% subsampling trials. **B-E** Connectivity differences of dementia in each subtype compared to all healthy control subjects. Wilcoxon rank-sum test was used to detect such differences. The top 50 (absolute) significant abnormal FCs of the first and second subtypes are visualized in panels **B** and **C.** Hyperconnections were represented in red, indicating that the FCs of healthy controls were larger than those in dementia patients for a subtype. Hypoconnections are represented in blue, indicating that FCs of healthy controls were smaller than those in dementia patients for a subtype. Node size indicated the node strength calculated from the sum of the absolute w value of linked FC. **D, E** Network-level FCs differences of dementia in subtype 1 and subtype 2, grouping from all ROI-level FCs significantly differed to HC.

To investigate the neurobiological basis of dementia patients within these subtypes, we examined the differences in FCs between dementia patients in each subtype and all healthy controls using the Wilcoxon rank sum test. We identified hypo-connections (significantly smaller FCs) and hyper-connections (significantly larger FCs) in dementia patients compared to healthy controls. In subtype 1, the largest hypo-connection was observed between the left insula (SMN) and left superior parietal cortex (DAN), while the largest hyper-connection was found between the right precentral gyrus (SMN) and right middle frontal cortex (FPC, Brodmann area 8) (Figure 4B, Figure S8A). The differences in brain connections mainly involved within-network connections of SMN and VAN, as well as between-network connections of SMN, FPC, and DAN (Figure 4D). In subtype 2, the largest hypo-connection was observed between the left inferior temporal cortex (DAN) and left inferior parietal cortex (DAN), while the largest hyper-connection was found between the left middle frontal cortex (DMN) and right supramarginal cortex (VAN) (Figure 4C, Figure S8B). The differences in brain connections mainly involved within-network connections of VIS and between-network connections of SMN, VIS, and DMN (Figure 4E). For subtype 3, no significant FC differences were observed compared to all healthy controls.

We further observed significant differences in the severity of NPS among dementia patients across the three subtypes, particularly in the hallucination, anxiety, and apathy domains (Table S5 and Figure S9). In addition, the anxiety, behavioral subsyndromes, and total NPS scores were significantly different between these subtypes. The patients in subtypes 1 and 2 exhibited more severe total NPS symptoms and subsyndromes compared to subtype 3 (Figures S9D-F). Moreover, functional impairment, as assessed by the FAQ and CDR, was more severe in patients of subtypes 1 and 2 in domains such as orientation, paying attention, and working on hobbies, in comparison to subtype 3 (Figure S9G, H, I). Notably, the presence of hallucination, anxiety, and apathy symptoms, as well as the two distinct neuropsychiatric subsyndromes, served as distinguishing factors between dementias in subtypes 1 and 2.

To further verify the uniqueness of the subtypes defined from NPS-linked FC latent features, we performed clustering directly using the NPI scores and compared the resulting clusters. As depicted in Figures S10A, B and C, two clusters were identified, which differed from the clusters derived using NPS-linked FC latent features (Figure S10D, with 41% of subjects assigned to consistent clusters). In these subtypes, the detected abnormal FCs in dementia were significantly decreased (Figure S8 and S10E, F). Additionally, functional disability, as evaluated by FAQ, was further enhanced and the differences in NPS symptoms were decreased (Tables S5 and S6), compared to the subtypes defined from FC-NPS linked latent features. In short, the NPS-FC linked features have an advantage in detecting the abnormal FCs and NPS difference compared to the neuropsychiatric symptoms.

### Longitudinal changes of characteristic phenotypes of dementia in three subtypes

Besides the baseline characteristic phenotypical difference, we revealed the longitudinal progression of dysfunction across various domains for different subtypes of dementia. Using a linear mixed-effects model to predict developmental trajectories of CDR, we observed different decline tendencies over time in orientation, hobbies maintenance, and general cognitive impairment for all three subtypes, as shown in Figure S11 A-E. Furthermore, we examined the longitudinal progression of cognitive functional change in animal naming, digits writing, and dots connecting tests measured by the NAB. The results revealed that patients in subtypes 1 and 2 exhibited similar rates of decline, which were both worse than those in subtype 3, as illustrated in Figure S11 I-K.

## DISCUSSION

Conducting sparse canonical correlation analysis between fMRI connectivity and NPS across subjects, we obtained two ten-fold cross-validated robust subsyndromes, behavioral and anxiety subsyndromes, in the NPS-FC correlated latent space. The behavioral and anxiety subsyndromes were associated with different connectivity patterns involving DMN, SMN and VIS as well as distinct characteristic phenotypes such as daily living disability. Then to comprehensively investigate the heterogeneity of dementia patients in relation to brain connectivity and NPS, we further divided all subjects into three subtypes based on the subsyndromes-linked FC latent features and analyzed the characteristic and neurobiological profiles of dementia patients within each subtype. Our results indicated significant differences in clinical measurements, baseline connectivity patterns, and longitudinal clinical progression among the three dementia subtypes. Notably, dementias in subtypes 1 and 2 exhibited more severe cognitive dysfunctions at baseline and a decline in cognitive abilities over time compared to healthy controls, while subtype 3 displayed similar brain and cognitive phenotypes to healthy controls. These findings underscore the potential for further research in developing more precisely targeted interventions tailored to the unique cognitive impairment and dysfunction observed in dementia of different subtypes.

### NPS subsyndromes and linked functional connectivity patterns

In our study, we found that the behavioral and the anxiety subsyndromes exhibited opposite associations with various clinical measurements such as the MMSE total score and sum of boxes of CDR in opposite directions. Notably, the overall pattern indicated the behavioral subsyndrome positively correlated with better cognitive performance, while the anxiety subsyndrome was positively associated with impaired cognitive performance. This may be due to the fact that, although these two subsyndromes were primarily dominated by different NPS, they partially overlapped in capturing neurobehavioral dimensions in an inverse manner. Specifically, the behavioral subsyndrome was negatively correlated with delusion and depression while the anxiety subsyndrome was positively correlated with hallucination and anxiety as shown in Figures 2B and 3B. Previous research has reported strong positive correlations between depression and anxiety (43), as well as between delusions and hallucinations (44). In addition, both subsyndromes were highly correlated with between-network connections involving VIS-SMN, DMN-SMN and DMN-VIS (Figures 2D and 3D). These findings are in line with studies that have demonstrated the presence of hallucinations and delusions in AD patients, which are associated with atrophy and decreased glucose metabolism of the visual network (45). Moreover, patients with AD or Parkinson’s disease have high metabolic activity in brain regions involving DMN and SMN, which positively correlates with anxiety and depression symptoms (46). Our current results, along with these previously reported findings, offer an explanation for the divergent relationship between the two subsyndromes and various cognition impairment measurements. Furthermore, these overlapping neuro-circuits and NPS symptoms captured by our subsyndromes align with the conceptual model, suggesting that cognitive impairment, accompanied by delusion, hallucination, anxiety and depression symptoms, is intricately related to DMN circuitry and atrophy of SMN and DMN (47).

Interestingly, our results revealed that the behavioral subsyndrome specifically captured variance in instrumental activities of daily living involving business affairs and driving function deficits. Patients requiring assistance in these activities had significantly lower behavioral subsyndrome scores. Such disabilities are known to be highly effective in discriminating between healthy individuals and those with mild dementia (48, 49). In addition, we found a significant correlation between the behavioral subsyndrome and connectivity within SMN network, which plays a key role in decision-making and driving ability (50, 51). This suggests that the behavioral subsyndrome might serve as a more sensitive psychopathological dimension in detecting mild cognitive functional impairment during the transition from healthy state to disease by specifically capturing decision-making deficits reflected by driving disability and SMN dysfunctions.

Furthermore, our study indicated a negative correlation between the anxiety subsyndrome and the fluency animal test as well as the delayed logical memory scores. This subsyndrome was primarily characterized by connections between DMN and VAN and connections within VIS. Previous studies have highlighted that verbal category fluency scores are correlated with the white matter hyperintensities in the frontal lobe (part of DMN) of dementia patients and activation of the bilateral lingual gyrus (part of VIS) in healthy controls (52, 53). Moreover, the decline in episodic memory performance as evaluated by the logical memory test has been related to altered DMN connectivity in patients with cognitive impairment (54). In line with these findings, the anxiety subsyndrome might be the psychopathological dimension that more specifically captured dysfunction in episodic memory and language processing, distinguishing it from the behavioral subsyndrome.

### Three dementia subtypes

Most of the existing studies on dementia subtypes have primarily focused on single modality including psychological or structural imaging or genetics data (55, 56). Remarkably, both abnormal functional connectome and neuropsychiatric symptoms have been identified as biomarkers for early dementia detection (57–59). But no prior studies have used neuropsychiatric symptom-linked connectivity features to uncover subtypes. In our study, we defined subtypes based on NPS-related FC latent features, which demonstrated considerable potential in disentangling the heterogeneity of neuropsychiatric and neurobiological information related to cognitive dysfunction of early dementia.

We observed shared connectivity differences in dementia of subtypes 1 and 2, compared to healthy controls, including connections within VIS and connections between VIS and SMN, between FPC and SMN, and between DMN and VAN. These network abnormalities were associated with deficits in social conduct, emotional processing and episodic memory retrieval. Consequently, dementia patients in subtypes 1 and 2 performed poorer in orientation and game, hobby maintenance than those in subtype 3 and healthy controls. Additionally, cognitive dysfunction deteriorated more rapidly in subtypes 1 and 2. Dysfunctions in brain networks such as DMN, FPC, SMN, and VAN have been associated with cognitive and behavioral abnormalities in various dementia types including Alzheimer’s disease, frontotemporal dementia and dementia with Lewy bodies (14, 60). Thus, these shared FC dysfunctions and cognitive abnormalities observed across our two dementia subtypes and the traditional dementia subtypes might be a common cognitive impairment-related neuropathological mechanism.

Remarkably, at baseline, dementia in subtype 2 had more severe symptoms of hallucination and anxiety than those in subtype 1. Increased severity of hallucination might be explained by notable differences of connectivity within VIS and between VIS and SMN, both observed in subtype 2, as hallucinations have been linked to pathological changes in the visual network (61). This visual dysfunction may contribute to other emotional control-related neuropsychiatric symptoms such as anxiety and apathy (61, 62). The enhanced difference of FCs involving DMN in subtype 2 compared to subtype 1, known for its role in emotion regulation may explain this contribution. Moreover, as age increased, dementia in subtype 2 showed more severe cognitive and behavioral dysfunction than dementia in subtype 1, assessed by CDR, NPS and NAB items. Previous research has suggested that visual impairment increases the risk of cognitive impairment and incident dementia (63). Based on our findings, it is plausible that the more severe hallucination of subtype 2 could be associated with the progression to dementia-related cognitive impairment, but more studies are needed to confirm the interaction between these factors and to explore the underlying mechanism supporting this hypothesis.

The dementia patient in subtype 3 was found to be similar to the healthy control group, with no significant difference in brain connectivity and presenting less severe NPS, subsyndromes, total score, and cognitive dysfunction in paying attention and orientation compared to subtypes 1 and 2. This aligns with the previous finding that some patients with brain disorders do not exhibit biological abnormalities and functional deficits compared to healthy controls (64). The presence of such heterogeneity in dementia underscores the importance for clinicians and researchers to consider these variations when developing more precise diagnostic tools and targeted interventions that align with the unique cognitive profiles and underlying neuropathological mechanisms of dementia patients.

There are also some limitations and potential extensions to be considered. First, other biological information such as genomics and proteome sequence also contribute to the heterogeneity of dementia. Future studies could explore these factors to gain a more comprehensive understanding of dementia subtypes and their underlying mechanisms. Second, the generalizability of the subsyndrome and subtypes defined in our study should be further validated in large and matched datasets. This validation would confirm whether the identified subsyndromes and subtypes accurately represent the broader dementia population and can be reliably applied in clinical settings. Third, the present study focused on the baseline participants for seeking subsyndrome and subtypes. Incorporating longitudinal imaging and clinical data in future research would provide a deeper understanding of the progressive nature of dementia and the interplay between brain-psychopathology dimensions over time.

These caveats notwithstanding, our study offers important insights into the heterogeneity of dementia under a data-driven analytical framework. We identified behavioral and anxiety subsyndromes that capture critical distinctions shared between brain circuits and NPS in dementia. Specifically, the behavioral subsyndrome linked-FC latent features were specifically and negatively associated with dysfunction in daily activities related to taxes affairs while the anxiety subsyndrome-linked FC latent features were specifically and positively linked to memory and language deficits. In addition, using these two subsyndromes, our extensive clustering analyses revealed three stable subtypes. The dementia patients in two subtypes exhibited significant differences in FC involving DMN, FPC, and SMN, which were associated with memory and emotional dysfunction both at baseline and in longitudinal progression. The third subtype resembled healthy controls in terms of cognitive function and functional connectivity. Overall, our study provides a unique perspective on the complex heterogeneity of dementias and underlying neurobiological mechanisms, which may ultimately inform the development of more targeted interventions and diagnostic tools for patients in different subtypes.

## Acknowledgement

This work was supported in part by Alzheimer’s Association Grant (AARG-22-972541), NIH grant nos. R01MH129694 and R21MH130956, and Lehigh University FIG (FIGAWD35), CORE, and Accelerator grants. Portions of this research were conducted on Lehigh University’s Research Computing infrastructure partially supported by NSF Award 2019035. GAF was supported by NIH grant nos. K23MH114023 and R01MH125886 and grants from the Brain and Behavior Research Foundation and One Mind – Baszucki Brain Research Fund.

## Supplementary Materials

### Supplementary results

**Figure S1.**
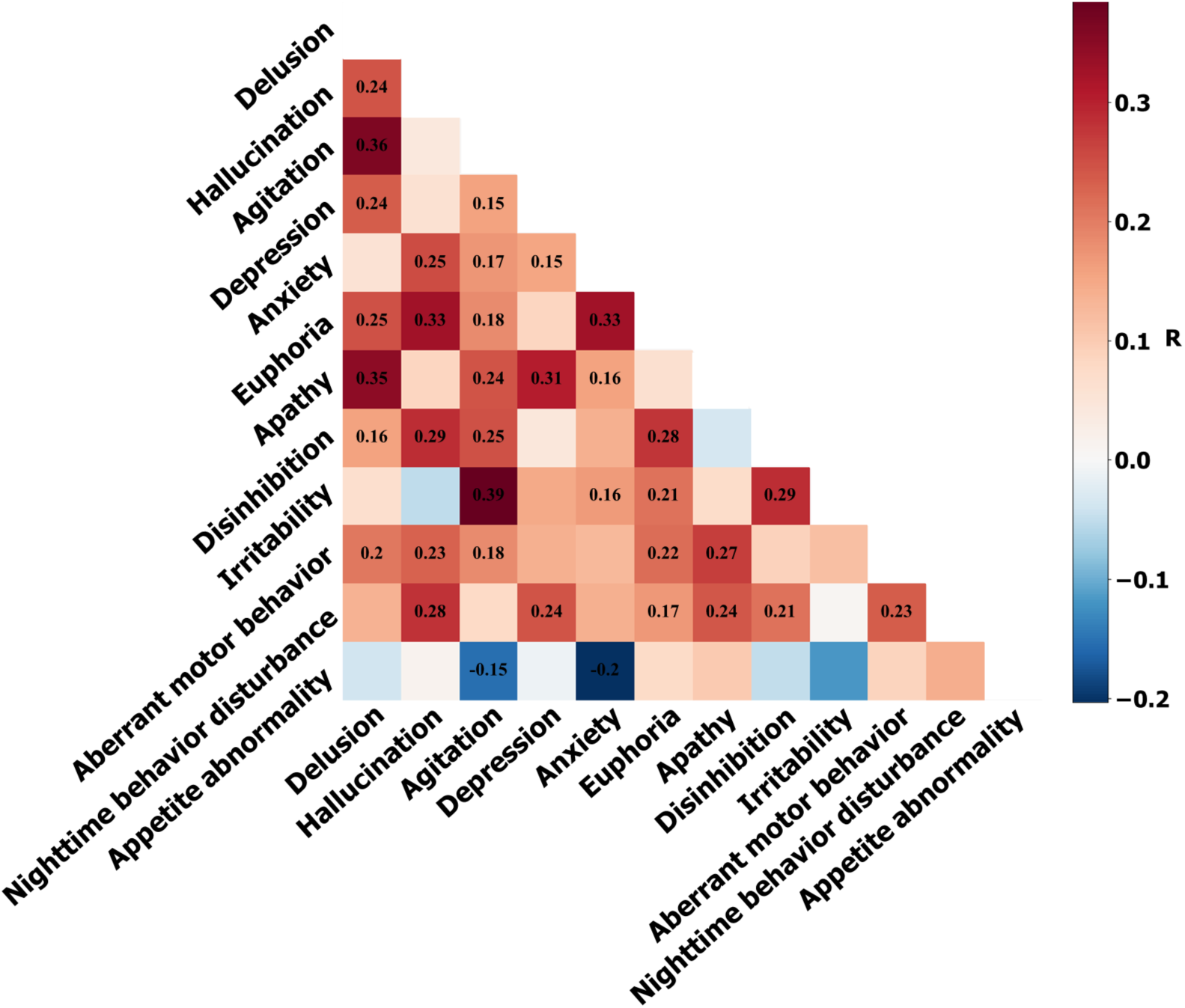
Correlation between pairs of neuropsychiatric symptoms. Only the significant correlations (p<0.05) were texted in the figure.

**Figure S2.**
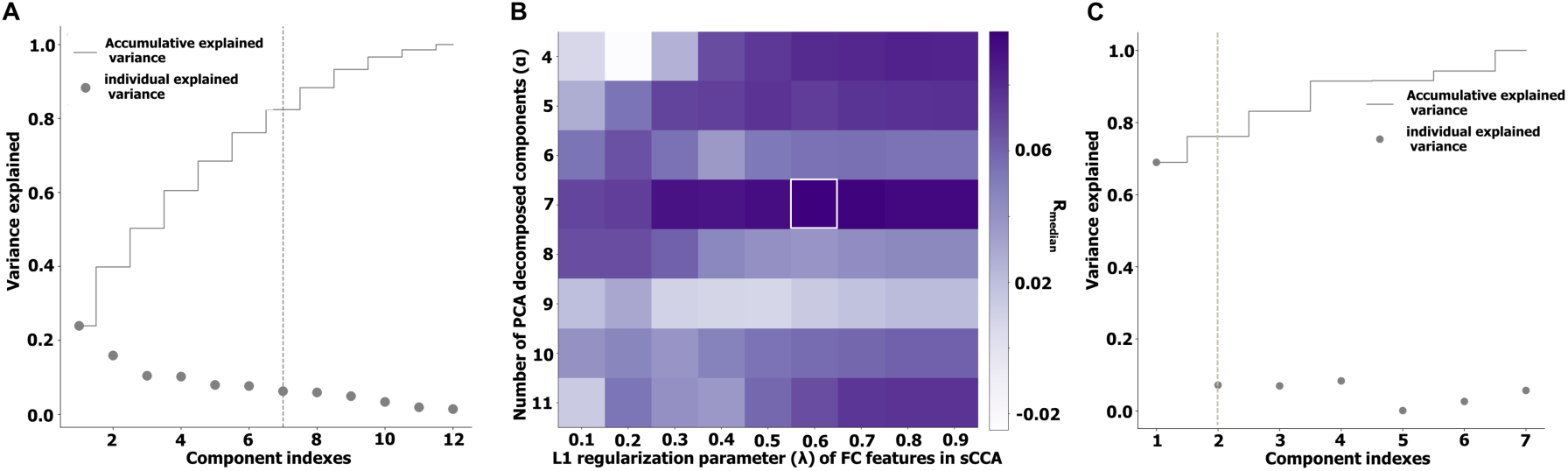
Hyperparameters selection in our study. **A** Variance explained by the principal components of NPS. **B** Average correlation of transformed pairwise canonical variates of ten-fold cross validation, with varying numbers of principal components of NPS and different L1 regularization parameters of connectivity features in sCCA. **C** Covariance explained by the canonical variates derived from sCCA between functional connectivity features and the first seven principal components of NPS.

**Figure S3.**
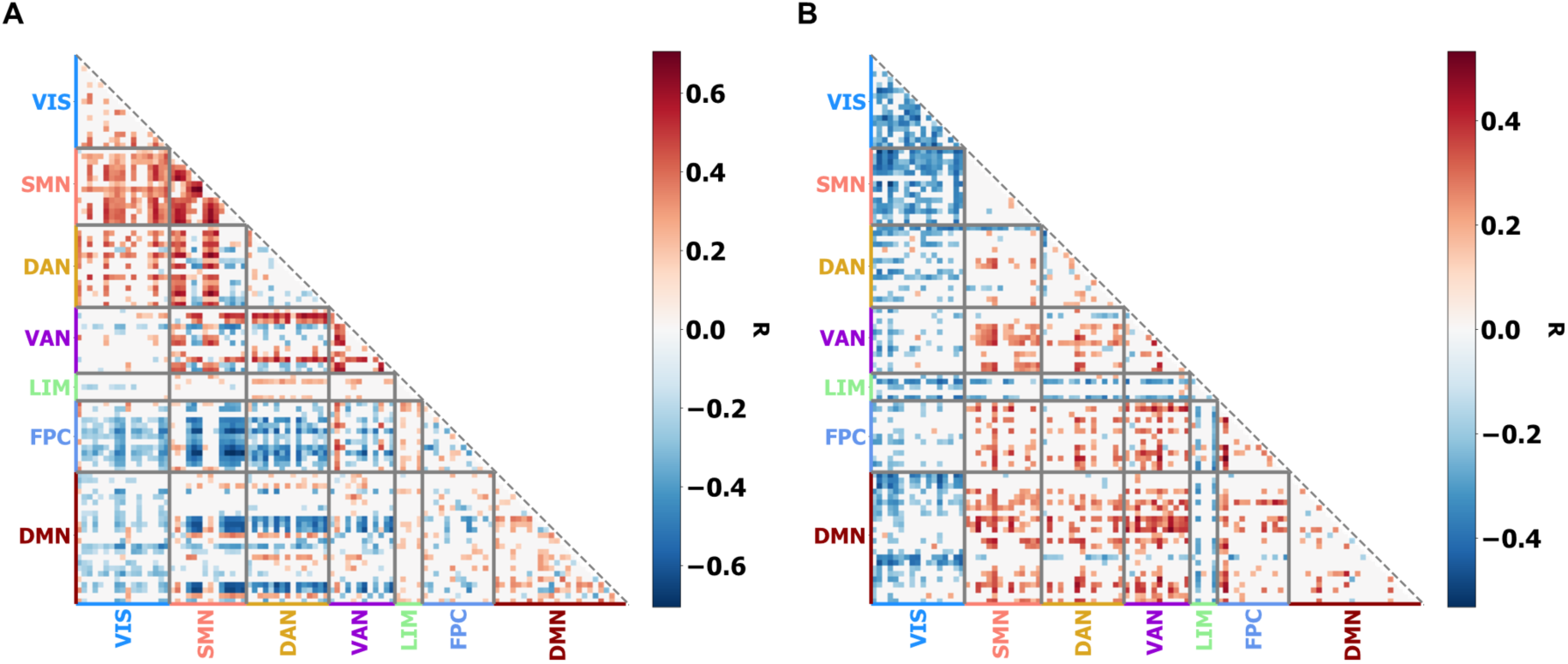
Significant correlations (after FDR correction) between FCs and the first and second sCCA transformed FC variates. Red color indicates positive correlation and blue color indicates negative correlation. (A) Correlation between FCs and the behavioral subsyndrome-linked FC latent features. (B) Correlation between FCs and the anxiety subsyndrome-linked FC latent features.

**Figure S4.**
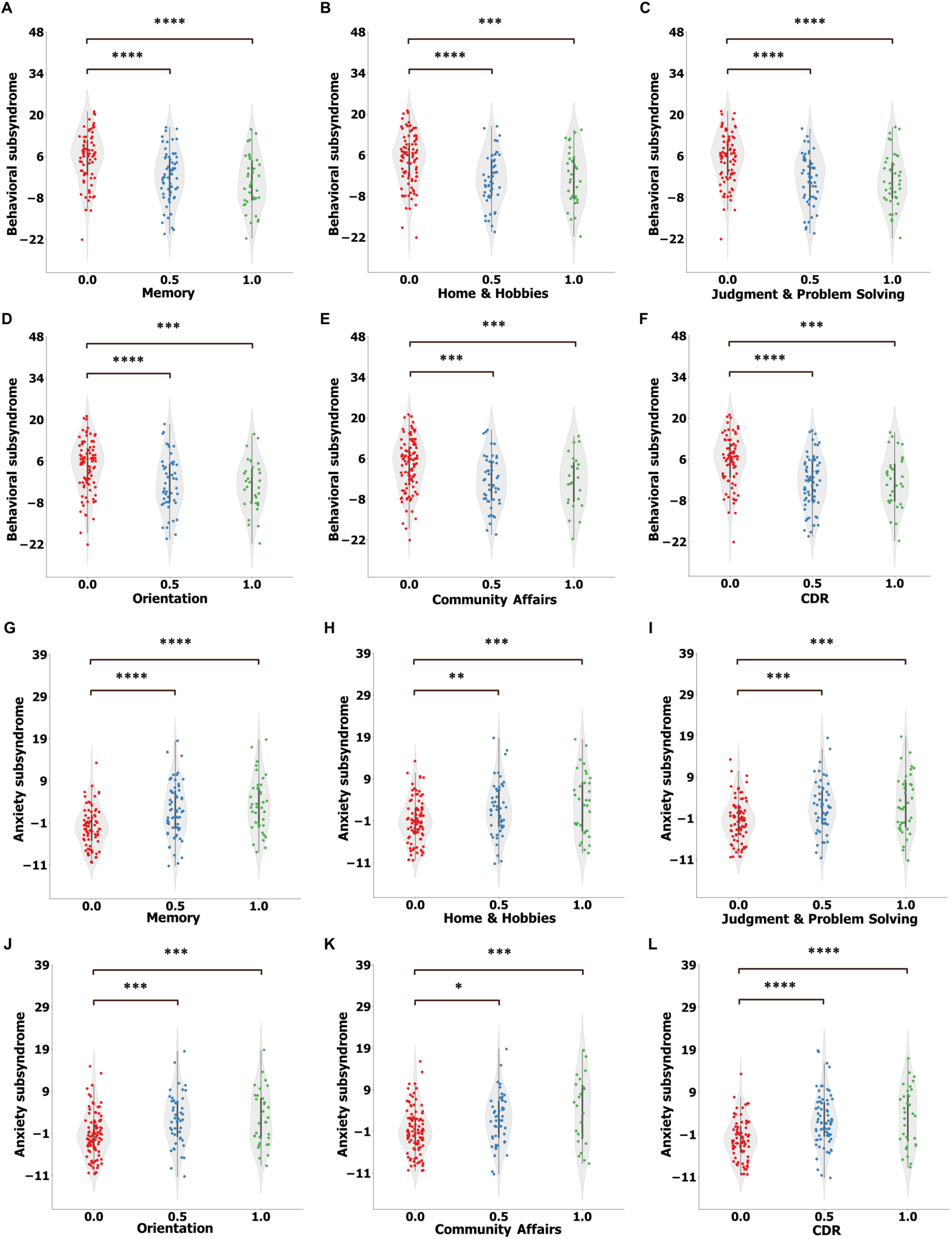
Tukey’s honest significance (HSD) test results for the behavioral (A-F) and anxiety (G-L) subsyndromes across groups with different CDR and its subscale scores. All p values were FDR corrected. (NS: p > 0.05; *: p ≤ 0.05; **: p ≤ 0.01; ***: p ≤ 0.001; ****: p ≤ 0.0001). The HSD test was not conducted for the personal care subscale as it was a binary categorical variable.

**Figure S5.**
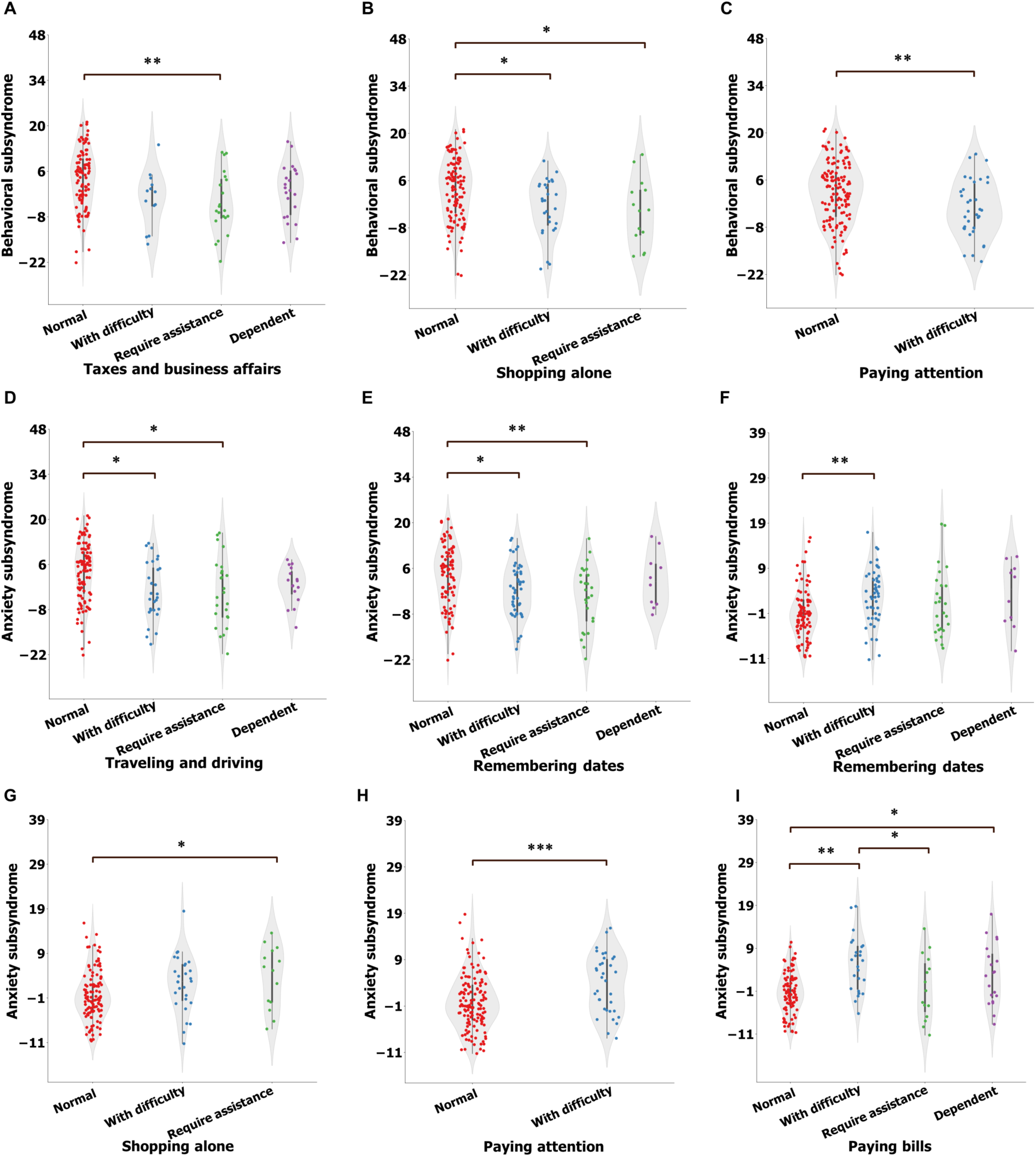
Tukey’s honest significance (HSD) test of the first (A-E) and second (F-I) canonical variates across groups with different functional assessment items. Pairwise difference analysis using HSD was conducted only for the items that showed significant differences in canonical variates across all groups, as determined by Kruskal-Wallis analysis. All p values were FDR corrected. (NS: p > 0.05; *: p ≤ 0.05; **: p ≤ 0.01; ***: p ≤ 0.001; ****: p ≤ 0.0001).

**Figure S6.**
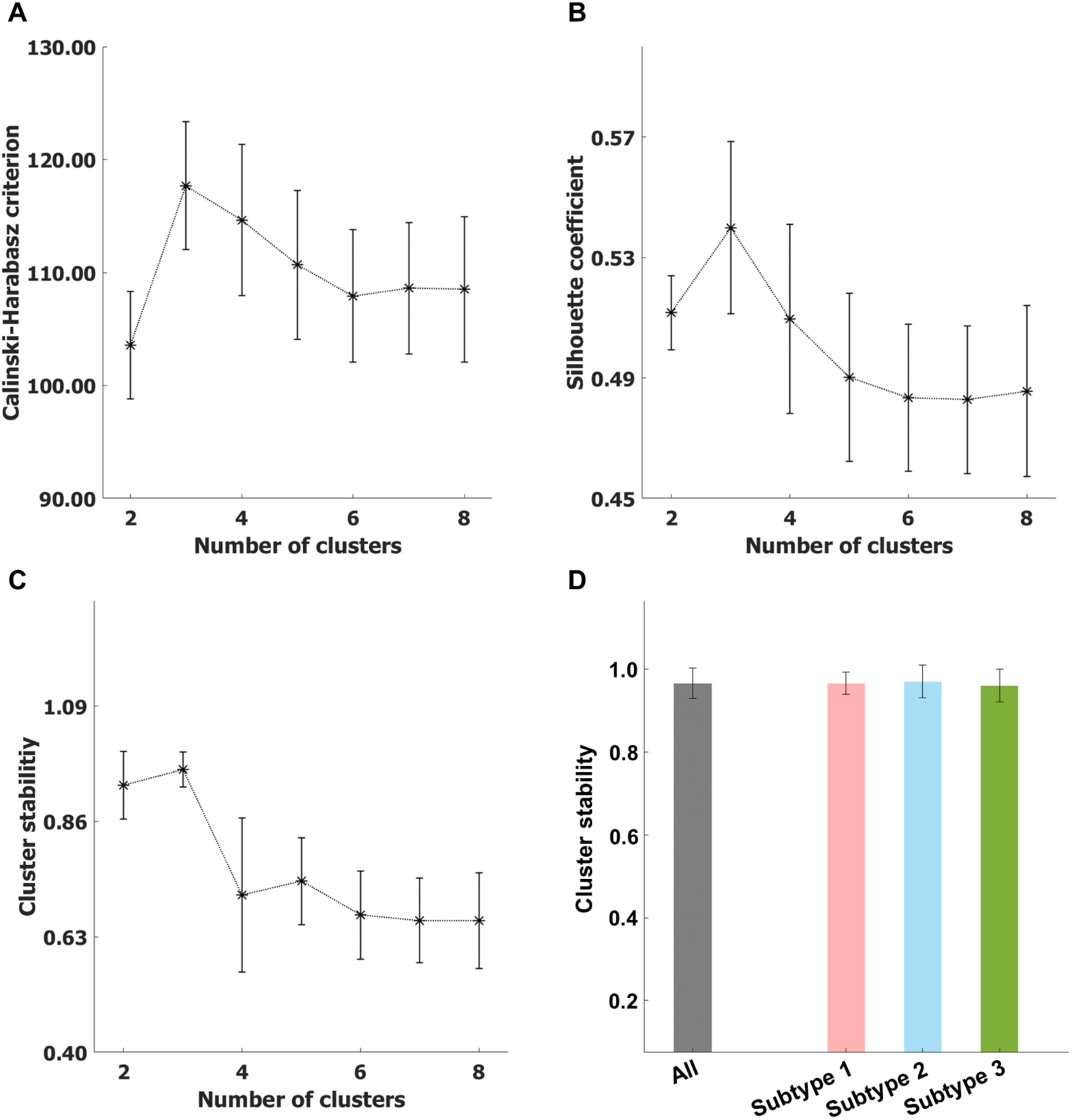
Cluster evaluation analyses of K-means clustering. We repeated K-means cluster 1000 times, with 90% random subsampling of all subjects. When cluster number (k) was 3, **A** the Calinski-Harabasz score and **B** Silhouette score, were maximized. **C** The stability coefficient, represented as the ratio of the same cluster label assigned to each subject across 1000 subsamplings, was calculated. The stability coefficients of different cluster numbers were plotted. **D** When k = 3, the stability coefficients of subjects in different subtypes were computed.

**Figure S7.**
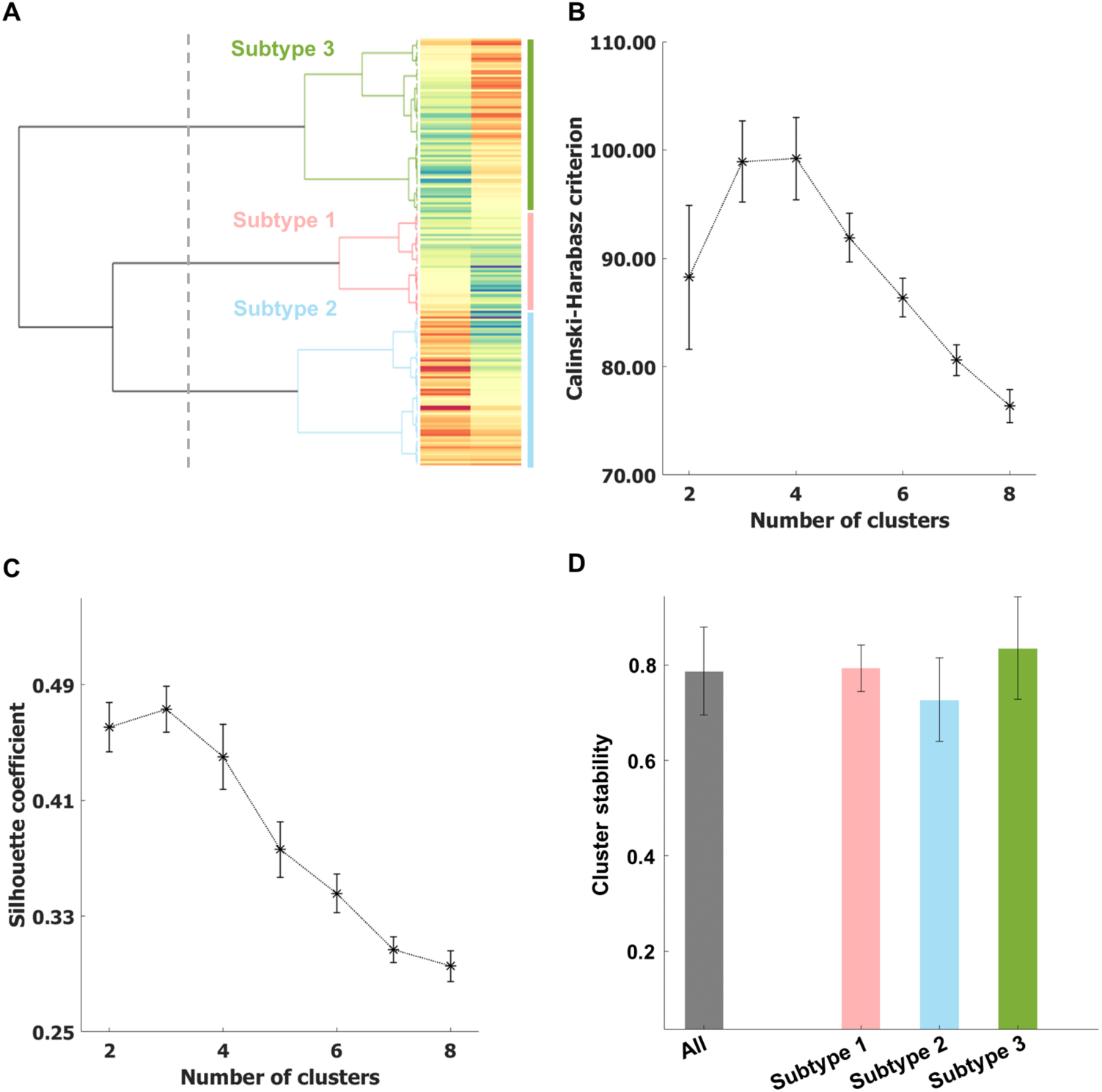
Cluster evaluation analyses of hierarchical clustering. **A** The visualization of hierarchical cluster result, when k = 3. We then repeated hierarchical clustering 1000 times, with 90% random subsampling of all subjects. When cluster number (k) was 3, **B** the Calinski-Harabasz score and **C** Silhouette score, were maximized. **D** To assess stability of the cluster analysis when cluster number was 3, the stability coefficient of different subtypes was computed as the ratio of the same cluster label assigned to subjects in each subtype across the 1000 subsamplings.

**Figure S8.**
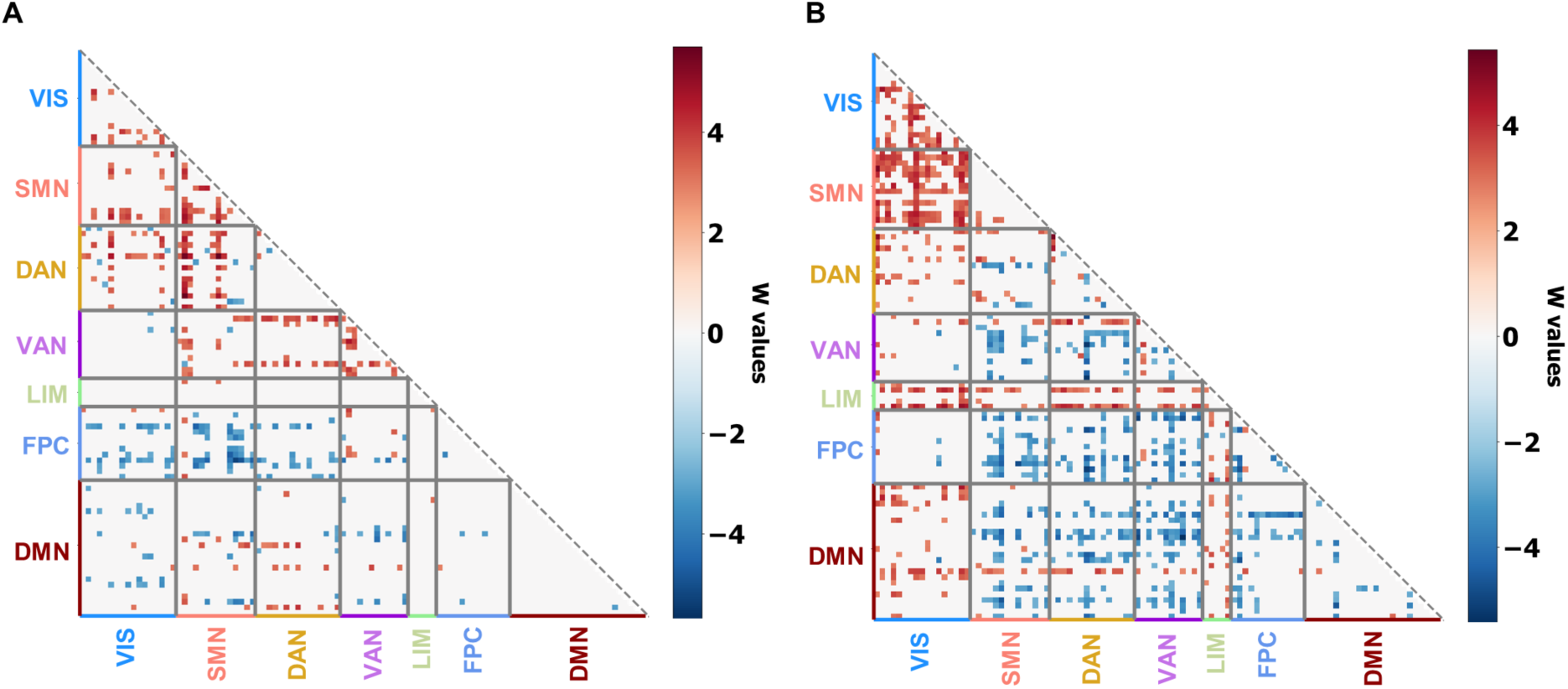
Differences of FCs between dementia of subtype 1 (A) and subtype 2 (B), compared to all healthy control subjects. The differences were detected using the Wilcoxon rank sum test and the significance of differences was corrected by FDR.

**Figure S9.**
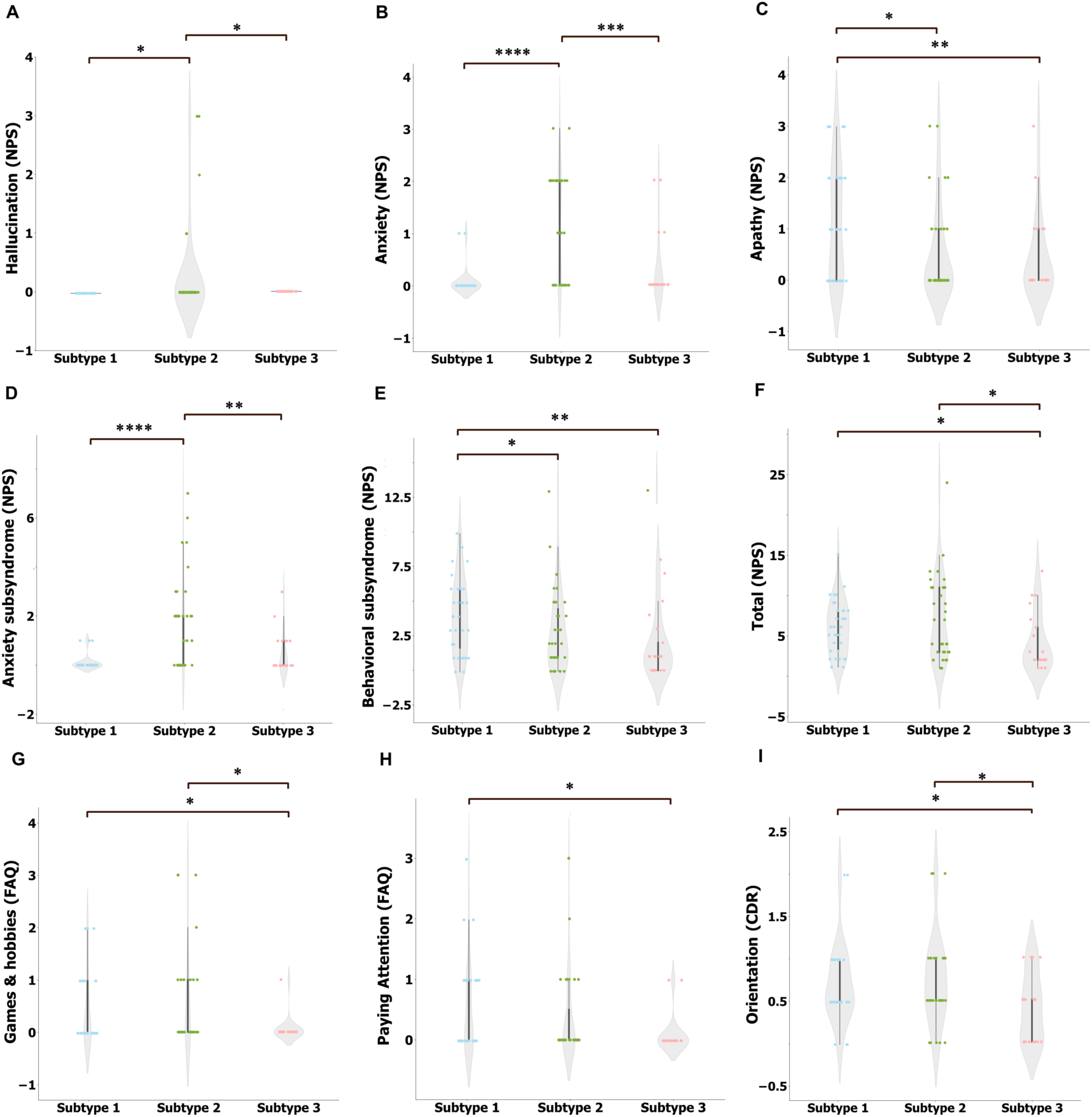
Dunn’s multiple comparison results of all ordinal clinical measurements, which were significantly different across three dementia subtypes. All p values were FDR corrected. (NS: p > 0.05; *: p ≤ 0.05; **: p ≤ 0.01; ***: p ≤ 0.001; ****: p ≤ 0.0001).

**Figure S10.**
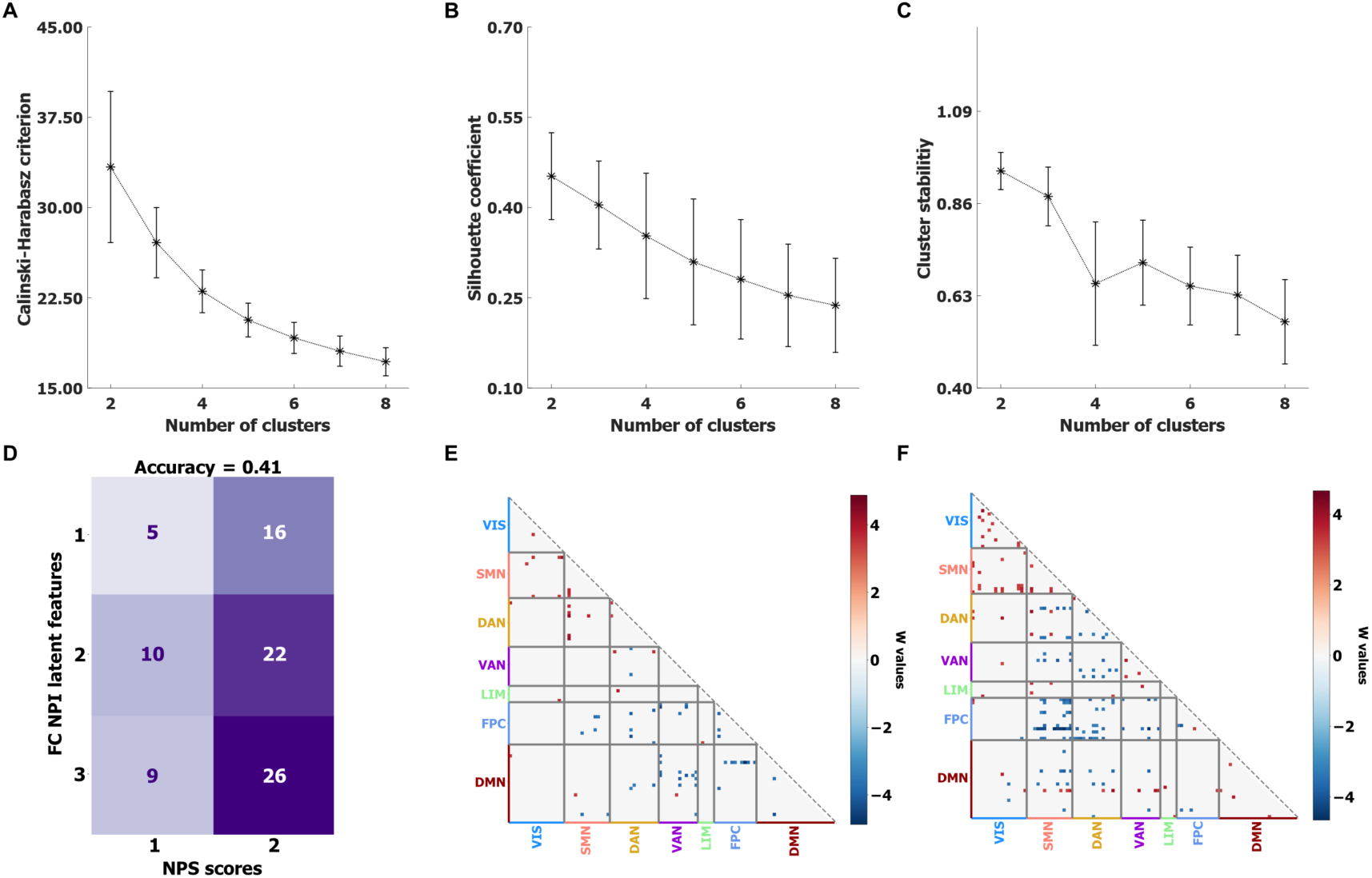
Clustering on NPS scores only. **A** Calinski-Harabasz score, **B** Silhouette score and **C** stability coefficient of different cluster numbers. **D** Confusion matrix comparing subtype labels obtained from clustering NPS scores only (x-axis) to clustering based on the NPS-linked FC latent scores (y-axis). The accuracy is 0.41. **E** Differences of FCs in dementia of subtype 1, compared to all healthy controls. **F** Difference of FCs in dementia of subtype 2, compared to all healthy controls. The differences were detected using the Wilcoxon rank sum test, and the significance of difference was corrected by FDR.

**Figure S11.**
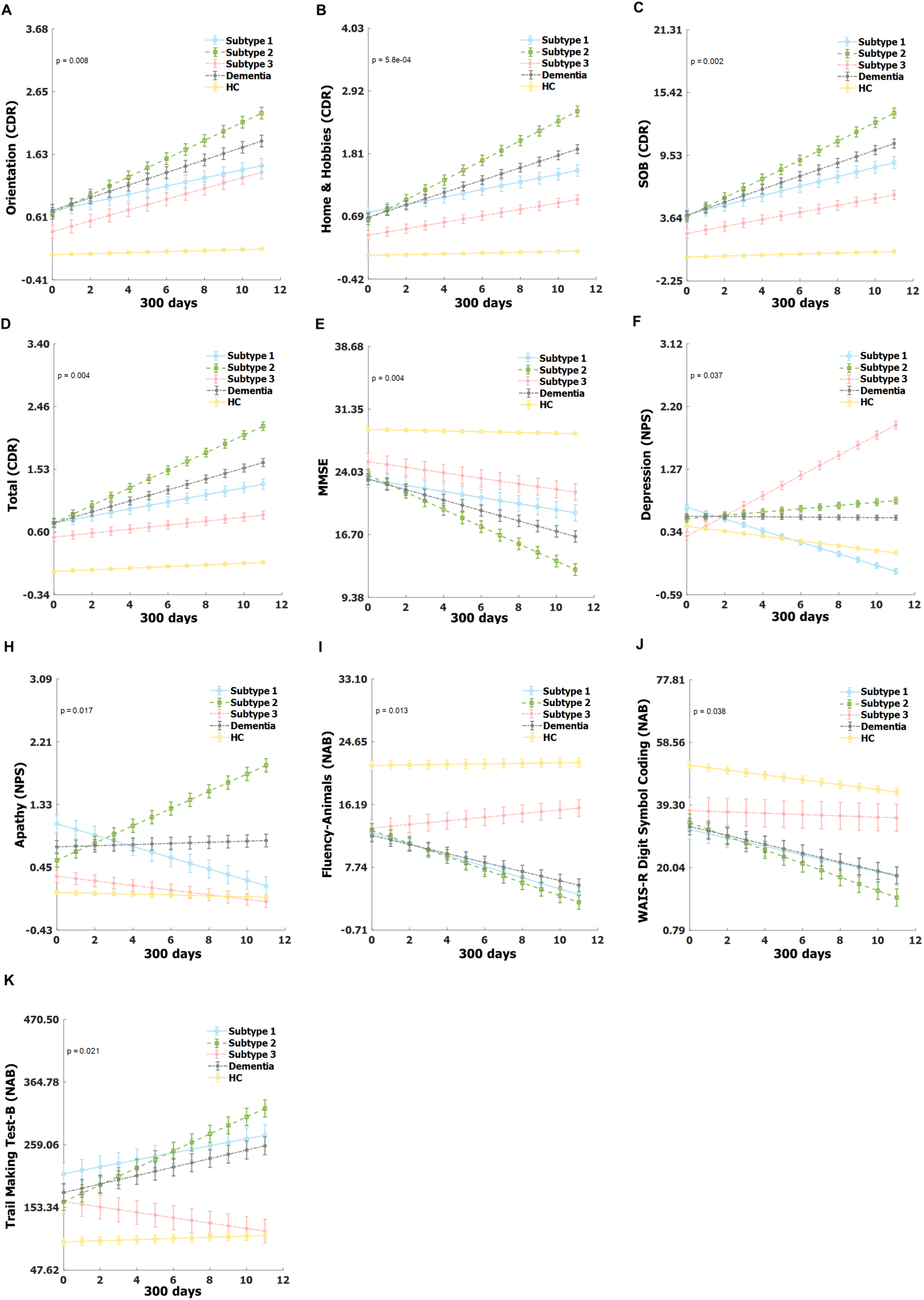
Differences in longitudinal change of various clinical scores among dementia in subtypes. Linear mixed effect models were employed to examine the difference in the longitudinal progression of clinical scores of dementia across subtypes. Only the clinical scores that exhibited significant differences in their longitudinal trajectories among the three dementia subtypes are presented.

**Table S1.** Summary the correlation between the third to seventh sCCA transformed connectivity latent scores and NPS latent scores. All p values were FDR corrected.

| **Canonical variates index** | **r** | **p** | **p<sub>permute</sub>** |
| --- | --- | --- | --- |
| 3<sup>rd</sup> | -0.174 | 0.020 | 0.974 |
| 4<sup>th</sup> | -0.013 | 0.859 | 0.583 |
| 5<sup>th</sup> | 0.072 | 0.337 | 0.459 |
| 6<sup>th</sup> | 0.161 | 0.033 | 0.201 |
| 7<sup>th</sup> | -0.006 | 0.941 | 0.583 |

**Table S2.**
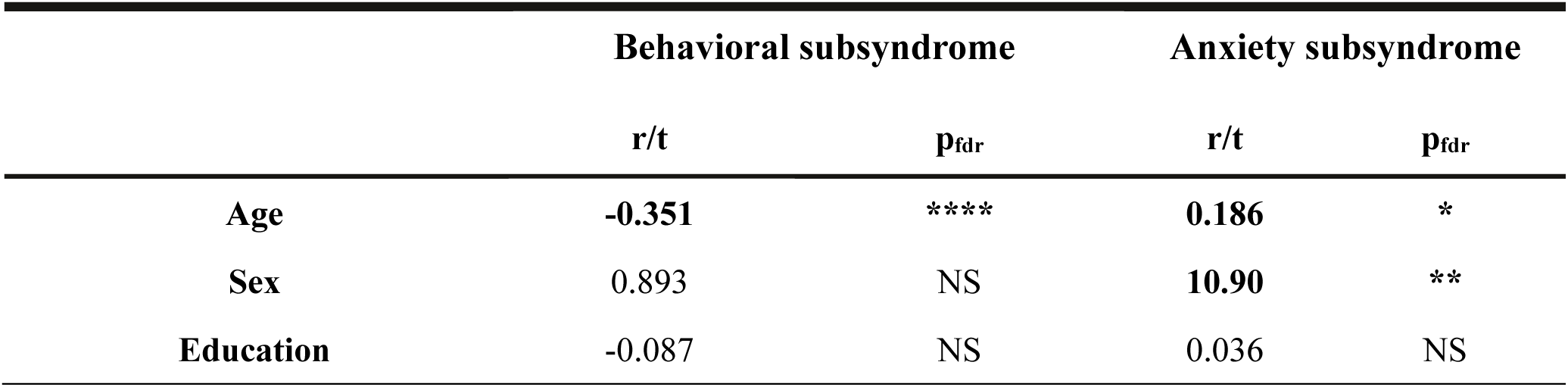
Associations between FC latent scores and demographic information. Two-sample t-test and Pearson correlation were applied. All p values were FDR corrected. (NS: p > 0.05; *: p ≤ 0.05; **: p ≤ 0.01; ***: p ≤ 0.001; ****: p ≤ 0.0001)

**Table S3.** Association between FC latent scores and the scores of various categorical clinical measurements and biomarkers. Statistical comparisons were performed using ANOVA and two-sample t-test. All p values were FDR corrected. (NS: p > 0.05; *: p ≤ 0.05; **: p ≤ 0.01; ***: p ≤ 0.001; ****: p ≤ 0.0001)

|  | Behavioral subsyndrome |  | Anxiety subsyndrome |  |
| --- | --- | --- | --- | --- |
|  | f/t | p<sub>fdr</sub> | f/t | p<sub>fdr</sub> |
| **Personal Care** | 2.26 | NS | 2.08 | NS |
| **Memory (CDR)** | **13.40** | **** | **13.50** | **** |
| **Home & hobbies (CDR)** | **10.20** | **** | **7.38** | *** |
| **Judgment & problem solving (CDR)** | **14.70** | **** | **7.94** | *** |
| **Orientation (CDR)** | **9.74** | **** | **8.89** | *** |
| **Community Affairs (CDR)** | **7.64** | *** | **7.44** | *** |
| **Total (CDR)** | **12.70** | **** | **13.50** | **** |
| **SOB (CDR)** | **-0.274** | *** | **0.345** | **** |
| **APOE** | 1.56 | NS | 2.18 | NS |
| **Diagnosis Label** | **4.95** | **** | **-4.41** | **** |
| **Paying Bills (FAQ)** | 2.33 | NS | **10.2** | **** |
| **Taxes and Business Affairs (FAQ)** | **5.27** | ** | 2.67 | NS |
| **Shopping Alone (FAQ)** | **5.88** | ** | **5.20** | * |
| **Games and Hobbies (FAQ)** | 4.65 | NS | 1.23 | NS |
| **Using Stove (FAQ)** | 2.31 | NS | 1.42 | NS |
| **Preparing a Balanced Meal (FAQ)** | 1.05 | NS | 2.07 | NS |
| **Current Events (FAQ)** | 1.82 | NS | 2.89 | NS |
| **Paying Attention (FAQ)** | **6.71** | * | **11.8** | ** |
| **Remembering Dates (FAQ)** | **4.70** | ** | **4.60** | ** |
| **Traveling and Driving (FAQ)** | **4.96** | ** | 2.83 | NS |

**Table S4.** Relationship between FC latent scores and the continuous clinical measurements was measured by using Pearson correlation. All p values were FDR corrected. (NS: p > 0.05; *: p ≤ 0.05; **: p ≤ 0.01; ***: p ≤ 0.001; ****: p ≤ 0.0001)

|  | Behavioral subsyndrome |  | Anxiety subsyndrome |  |
| --- | --- | --- | --- | --- |
|  | r | p<sub>fdr</sub> | r | p<sub>fdr</sub> |
| **MMSE** | **0.309** | **\*\*\*\*** | **-0.304** | **\*\*\*\*** |
| **Digit Span-Forward (NAB)** | 0.135 | NS | 0.002 | NS |
| **Digit Span-Backward (NAB)** | 0.081 | NS | -0.087 | NS |
| **Fluency-Animals (NAB)** | 0.129 | NS | **-0.232** | **\*\*** |
| **Fluency-Vegetable (NAB)** | **0.243** | **\*\*** | **-0.255** | **\*\*\*** |
| **Trail Making Test-A (NAB)** | **-0.190** | **\*** | **0.222** | **\*\*** |
| **Trail Making Test-B (NAB)** | -0.189 | NS | **0.221** | **\*** |
| **WAIS-R Digit Symbol Coding (NAB)** | **0.224** | **\*\*** | **-0.262** | **\*\*\*** |
| **Logical Memory Immediate (NAB)** | **0.201** | **\*** | **-0.232** | **\*\*** |
| **Logical Memory Delayed (NAB)** | 0.159 | NS | **-0.226** | **\*\*** |
| **Boston (NAB)** | **0.193** | **\*** | **-0.168** | **\*** |

**Table S5.**
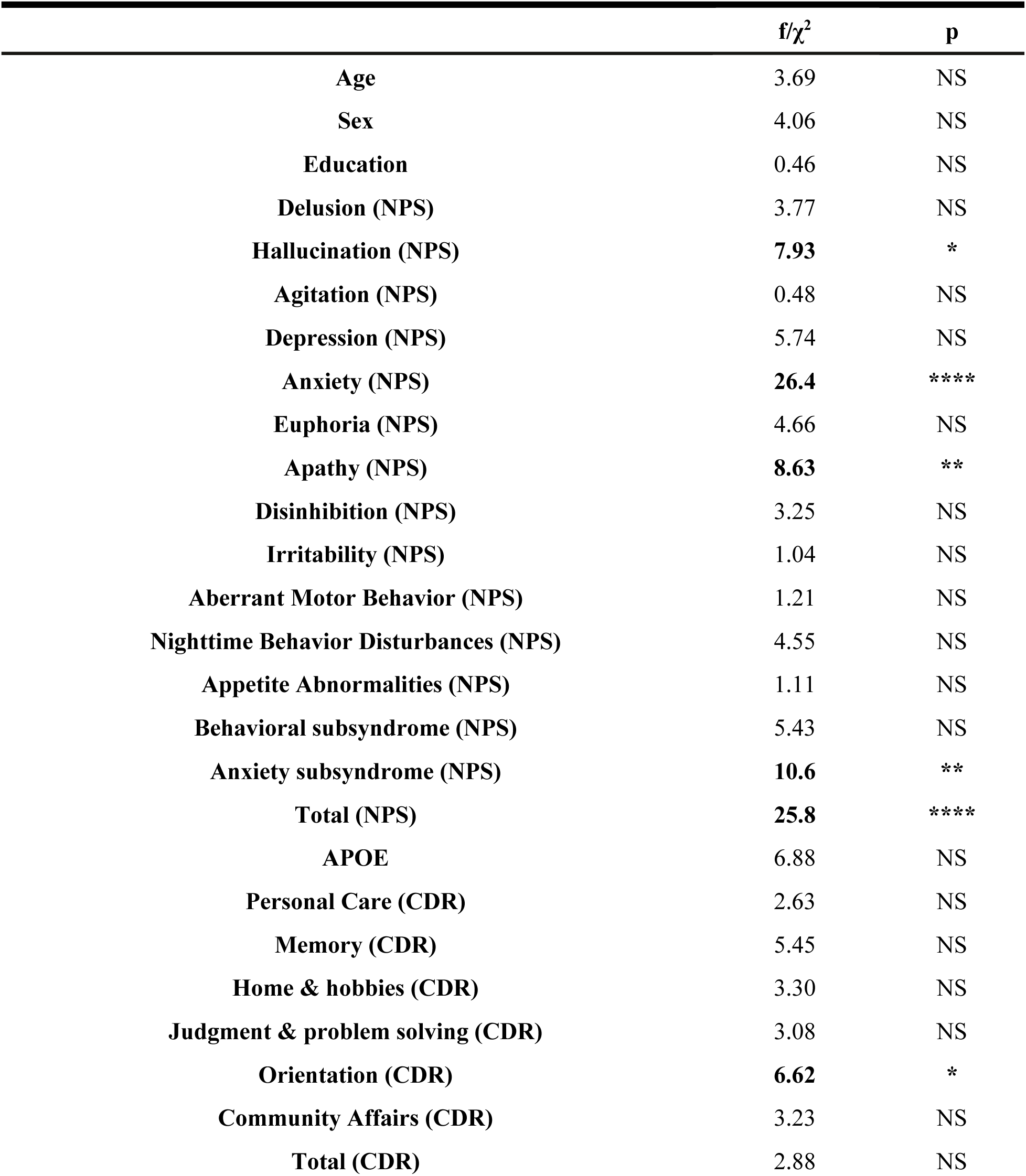
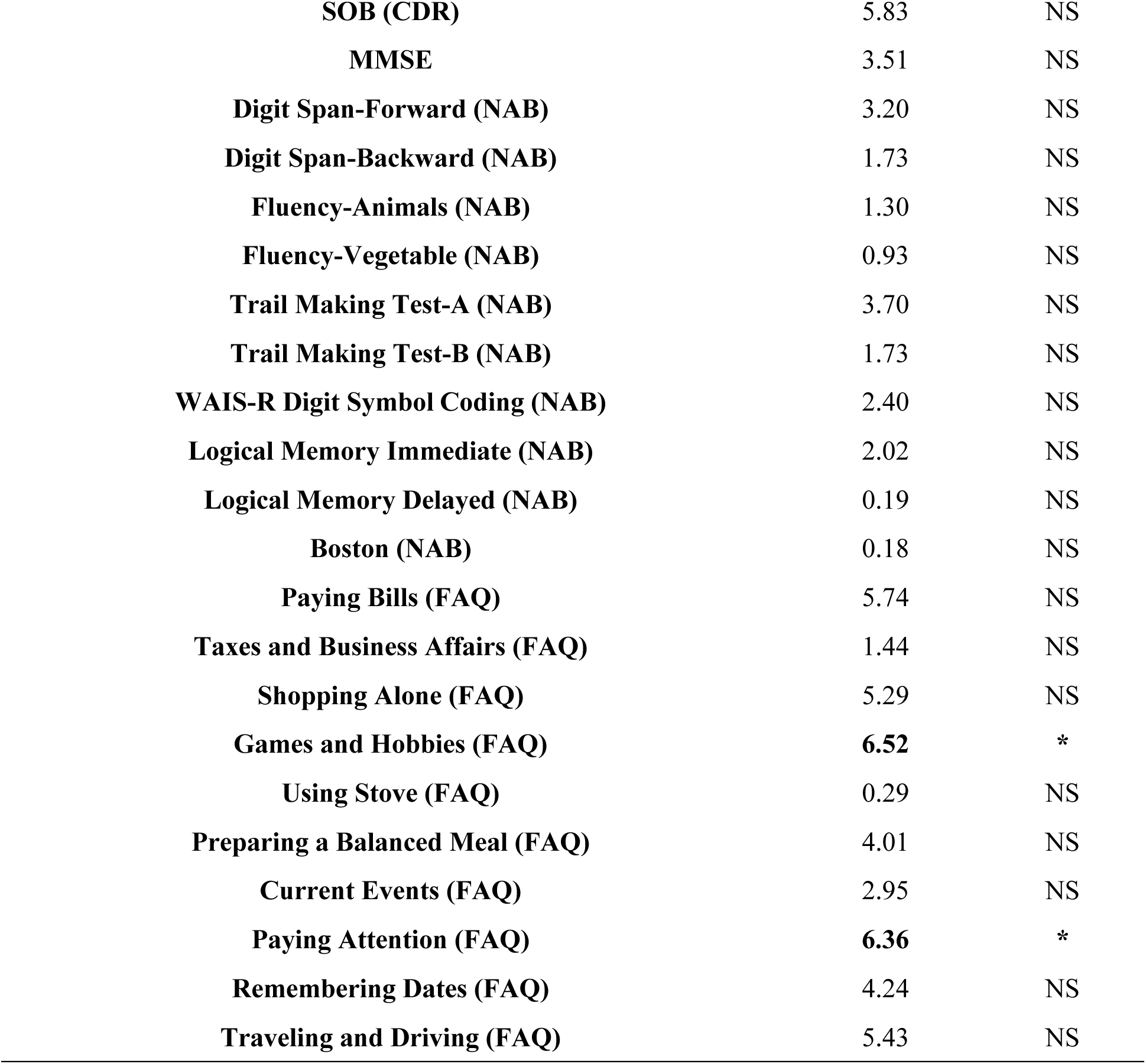
Differences of various clinical measurements and demographic information across dementia in three subtypes defined from FC-NPS linked latent features. Kruskal–Wallis ANOVA and Chi-square were applied in statistic comparisons of ordinal and categorical variables respectively, unless corresponding variables of each group were equal. (NS: p > 0.05; *: p ≤ 0.05; **: p ≤ 0.01; ***: p ≤ 0.001; ****: p ≤ 0.0001)

**Table S6.** Difference of various clinical measurements and demographic information across dementia in three subtypes defined from NPS. Kruskal–Wallis ANOVA and Chi-square were applied in the statistic comparisons of ordinal and categorical variables respectively, unless corresponding variables of each group were equal. (NS: p > 0.05; *: p ≤ 0.05; **: p ≤ 0.01; ***: p ≤ 0.001; ****: p ≤ 0.0001)

| | $f/\chi^2$ | p |
| --- | --- | --- |
| **Age** | 0.09 | NS |
| **Sex** | 5.89 | NS |
| **Education** | 2.07 | NS |
| **Delusion (NPS)** | 3.39 | NS |
| **Hallucination (NPS)** | 1.43 | NS |
| **Agitation (NPS)** | 3.87 | NS |
| **Depression (NPS)** | 0.89 | NS |
| **Anxiety (NPS)** | 4.97 | NS |
| **Euphoria (NPS)** | 5.90 | NS |
| **Apathy (NPS)** | 5.24 | NS |
| **Disinhibition (NPS)** | 0.32 | NS |
| **Irritability (NPS)** | 1.31 | NS |
| **Aberrant Motor Behavior (NPS)** | 5.96 | NS |
| **Nighttime Behavior Disturbances (NPS)** | 1.50 | NS |
| **Appetite Abnormalities (NPS)** | 2.32 | NS |
| **Behavioral subsyndrome (NPS)** | 3.07 | NS |
| **Anxiety subsyndrome (NPS)** | **6.08** | * |
| **Total (NPS)** | 3.46 | NS |
| **APOE** | 3.89 | NS |
| **Personal Care (CDR)** | 2.40 | NS |
| **Memory (CDR)** | 1.36 | NS |
| **Home & hobbies (CDR)** | 1.36 | NS |
| **Judgment & problem solving (CDR)** | 0.96 | NS |
| **Orientation (CDR)** | 1.72 | NS |
| **Community Affairs (CDR)** | 2.26 | NS |
| **Total (CDR)** | 1.37 | NS |
| **SOB (CDR)** | 1.62 | NS |
| **MMSE** | 0.14 | NS |
| **Digit Span-Forward (NAB)** | 3.63 | NS |
| **Digit Span-Backward (NAB)** | 1.41 | NS |
| **Fluency-Animals (NAB)** | 0.38 | NS |
| **Fluency-Vegetable (NAB)** | 1.26 | NS |
| **Trail Making Test-A (NAB)** | 0.13 | NS |
| **Trail Making Test-B (NAB)** | 1.06 | NS |
| **WAIS-R Digit Symbol Coding (NAB)** | 0.97 | NS |
| **Logical Memory Immediate (NAB)** | 1.86 | NS |
| **Logical Memory Delayed (NAB)** | 3.23 | NS |
| **Boston (NAB)** | 0.82 | NS |
| **Paying Bills (FAQ)** | **12.7** | **\*\*** |
| **Taxes and Business Affairs (FAQ)** | **12.0** | **\*\*** |
| **Shopping Alone (FAQ)** | **13.2** | **\*\*** |
| **Games and Hobbies (FAQ)** | **14.8** | **\*\*\*** |
| **Using Stove (FAQ)** | **8.66** | **\*** |
| **Preparing a Balanced Meal (FAQ)** | 5.46 | NS |
| **Current Events (FAQ)** | **11.0** | **\*\*** |
| **Paying Attention (FAQ)** | **12.7** | **\*\*** |
| **Remembering Dates (FAQ)** | **16.5** | **\*\*\*** |
| **Traveling and Driving (FAQ)** | **16.5** | **\*\*\*** |

